# IFT cargo and motors associate sequentially with IFT trains to enter cilia

**DOI:** 10.1101/2023.06.20.545804

**Authors:** Aniruddha Mitra, Elizaveta Loseva, Erwin J.G. Peterman

## Abstract

Intraflagellar transport (IFT) orchestrates entry of proteins into primary cilia. At the ciliary base, assembled IFT trains, driven by kinesin-2 motors, can transport cargo proteins into the cilium, across the crowded transition zone (TZ). How trains assemble at the base and how proteins associate with them is far from understood. Here, we use single-molecule imaging in the cilia of *C. elegans* chemosensory neurons to directly visualize the entry of kinesin-2 motors kinesin-II and OSM-3, as well as anterograde cargo proteins IFT dynein and tubulin. Single-particle tracking shows that IFT components associate with trains sequentially, both in time and space. Super-resolution maps of IFT components in wild-type and mutant worms reveal ciliary ultrastructure and show that kinesin-II is essential for axonemal organization. Finally, imaging cilia lacking kinesin-II and/or TZ function uncovers the interplay of kinesin-II and OSM-3 in driving efficient transport of IFT trains across the TZ.

## Introduction

Primary or sensory cilia are organelles that protrude from most eukaryotic cells and are involved in detecting and relaying external cues to the cell body (Nachury and Mick, 2019). Proper ciliary function requires a tightly regulated pool of proteins in the cilioplasm and ciliary membrane, different in composition from the rest of the cell body. To maintain this heterogeneity, eukaryotic cells have evolved a specialized intracellular transport system called intraflagellar transport (IFT; Figure 1A) (Mul et al., 2022; Prevo et al., 2017). Mutations in IFT-associated proteins can disrupt the protein pool in the cilium, resulting in ciliopathies (Reiter and Leroux, 2017). Ciliary architecture, IFT-train structure, and IFT dynamics have been studied in detail in recent years. The core structural element is an axoneme, consisting of nine doublet microtubules (MTs) in a cylindrical configuration, with the doublets emanating from the basal body at the ciliary base (degenerated in *C. elegans*). Located further along the cilium is the transition zone (TZ), characterized by Y-shaped protein complexes that link the axoneme to the ciliary membrane, forming a dense region (Figure 1B) that acts as a diffusion barrier to proteins (Garcia-Gonzalo and Reiter, 2017), with most ciliary proteins capable of crossing the TZ only by hitching a ride on anterograde IFT trains. IFT trains are large (>80 MDa), ordered, polymeric structures with periodic repeats of IFT-B and IFT-A complexes (Jordan et al., 2018; Jordan and Pigino, 2021; Webb et al., 2020). Anterograde kinesin-2 motors associate with IFT-B proteins and drive the trains from the ciliary base, through the TZ, to the ciliary tip (Mul et al., 2022). In *C. elegans*, anterograde IFT is driven by two distinct kinesin-2 motors, heterotrimeric kinesin-II and homodimeric OSM-3. The slower kinesin-II is understood to navigate anterograde IFT trains across the TZ, after which it is gradually replaced at the ‘handover’ zone by the faster OSM-3 (Prevo et al., 2015), which drives the trains to the ciliary tip (Ou et al., 2005; Pan et al., 2006). At the ciliary tip, anterograde trains are reconfigured into less-structured retrograde trains (Jordan et al., 2018). IFT-dynein, autoinhibited cargo of anterograde IFT trains, drive retrograde trains from the ciliary tip to the ciliary base (Toropova et al., 2017).

**Figure 1:**
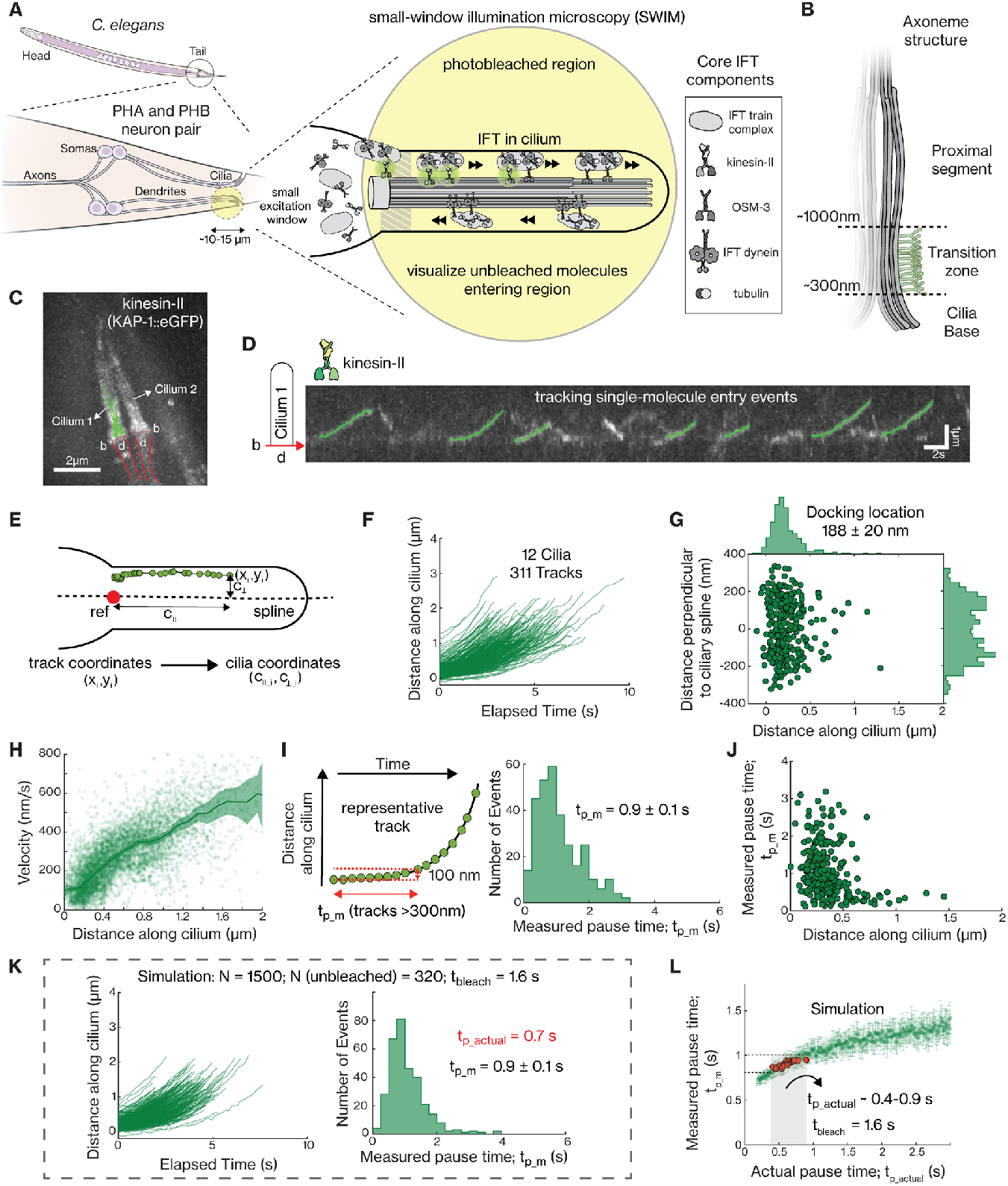
Analysis of the dynamics of kinesin-II (KAP-1::eGFP) molecules entering the cilia of PHA/PHB neurons. **(A)** Illustration of single-molecule imaging of kinesin-II motors entering the sensory cilia of PHA/PHB neurons located in the tail of *C. elegans*, using small-window illumination microscopy (SWIM). Using SWIM, we can visualize not-yet-photobleached kinesin-II molecules in a small excitation region for a long duration of time. **(B)** Illustration of the axonemal structure at the initial part of the cilia. The axoneme starts at the ciliary base, passes through the transition zone between ∼300-1000 nm, moving into the proximal segment. **(C)** Maximum projection of eGFP-labelled kinesin-II (KAP-1::eGFP) imaged in PHA/PHB cilia of a worm (see Supplementary Movie 2). Tracks of single-molecule events entering cilium 1 have been indicated (green lines), b indicates the base of the two cilia (dotted red lines) and d indicates the region where the corresponding dendrites are located (between the solid red lines). **(D)** Representative kymograph (space-time intensity plot) of kinesin-II along cilium 1, in 1C, display that single kinesin-II molecules, diffusing in the dendrite, dock at the ciliary base and enter the cilium. Single-molecule entry events are tracked (tracks indicated in green). **(E)** By drawing a central spline along the cilium and defining a reference point (at the ciliary base), x-y coordinates of tracks are transformed to ciliary coordinates, as shown in the scheme (also see Supplementary Figure 2). **(F)** Distance-time plots of all kinesin-2 tracks pooled together (311 tracks from 12 cilia). **(G)** Docking location of all kinesin-2 tracks at the ciliary base. Distributions of the docking location along the ciliary length (above the plot) and perpendicular to the ciliary spline (right of the plot). Kinesin-II molecules mostly dock between 0-1μm from the ciliary base with the average docking location at 289 ± 19 nm. **(H)** Distribution of velocity along the cilium length (6693 datapoints). Solid line represents the binned average velocity and shaded area indicates the error. **(I)** Left panel: Measured pause time, *t*_*p*_*m*_ is defined as the amount of time it takes for a track to move 100 nm, with only tracks longer than 300 nm included in the analysis. Right panel: Histogram of measured pause time (average pause time is 0.9 ± 0.1 s). **(J)** Correlation of *t*_*p*_*m*_ with docking location of kinesin-II events. **(K)** Distance-time plots of simulated kinesin-II entry events (left panel) and histogram of measured pause time (average pause time *t*_*p*_*m*_ = 0.9 ± 0.1 s; right panel), with *t*_*p*_*actual*_ = 0.7 s and *t*_*bleach*_ = 1.6 s (as calculated in Supplementary Figure 3B). Out of the 1500 simulated events (N) only 320 do not bleach before they move 300 nm from docked location. Simulation details are provided in Supplementary Figure 3. **(L)** Distribution of the measured pause times, *t*_*p*_*m*_, with respect to actual pause times, *t*_*p*_*actual*_, obtained from numerical simulations of kinesin-II molecules entering cilia, assuming a characteristic bleach time (*t*_*bleach*_) of 1.6 s. Each point represents the average pause time (*t*_*p*_*m*_) for a given simulated experiment (number of events without photobleaching N =400). *t*_*p*_*actual*_ is in the range 0.4-0.9 s for *t*_*p*_*m*_ 0.9 ± 0.1 nm (obtained from experiments; Figure 1I). Average value and error are estimated using bootstrapping (see Methods).

While recent studies have revealed key aspects of how IFT takes place inside cilia, the overall picture on the dynamic processes occurring at the ciliary base is still relatively incomplete. In fluorescence microscopy studies, we observed that the pools of IFT train, motor and cargo proteins are highly concentrated around the ciliary base, where anterograde IFT trains are assembled (Mijalkovic et al., 2017; Prevo et al., 2015). Careful FRAP (fluorescence recovery after bleaching) experiments have revealed that IFT proteins linger at the ciliary base for different durations before entering the cilium, suggesting slow, stepwise assembly of IFT trains (Hibbard et al., 2021; Wingfield et al., 2017). A recent structural study in *C. reinhardtii*, has shown that anterograde IFT trains are assembled in a sequential manner at the ciliary base, tethered at one end to the TZ (van den Hoek et al., 2022). It appears that IFT-B complexes form a template scaffold, with IFT-A, IFT-dynein complexes and kinesin-II binding at subsequent stages. Few studies have also explored how IFT cargoes cross the TZ to enter the cilia. Several ciliary membrane proteins have been shown to couple to IFT trains that move them across the TZ in a directed manner (van Krugten et al., 2022; Weiss et al., 2019; Ye et al., 2013). Crucial cytosolic ciliary proteins, such as IFT dynein and tubulin, also bind to anterograde IFT trains as cargo. IFT-dynein binds in an autoinhibited conformation, with neighboring IFT dyneins tightly packed, repeating every 18 nm (Toropova et al., 2019; Vuolo et al., 2020). Tubulin is known to bind anterograde IFT trains through several interactions with IFT-B proteins (Bhogaraju et al., 2013), but has also been shown to enter the cilium diffusively in *C. reinhardtii* (Craft et al., 2015; Craft Van De Weghe et al., 2020). While the structural features of IFT trains at the ciliary base have become clearer, the dynamics of how different ciliary proteins associate with IFT trains and navigate the TZ to enter the cilium have not yet been resolved.

In this study, we employ *C. elegans* chemosensory cilia as a model system to directly visualize how different IFT motors and cargo proteins enter the cilium. Chemosensory neurons of *C. elegans*, like the PHA/PHB neuron pairs located in the tail of the worms (Figure 1A), have a cilium separated from the cell soma by a long dendrite. Ciliary proteins, like IFT motors and tubulin, form a small dynamic pool of diffusive proteins at the periciliary membrane compartment (PCMC; located at the transition between dendrite and cilium), near the ciliary base. Here, we utilize this geometry of *C. elegans* chemosensory neurons to perform single-molecule fluorescence imaging at the ciliary base, by exciting (and photobleaching) only a small section of the neuron occupied by the PCMC and the cilia (∼10 μm; Figure 1A). Using small-window illumination microscopy (SWIM) (Mitra et al., 2022), we can continuously image single fluorescent molecules entering the small illuminated region for a relatively long duration of time, since the diffusive pool of proteins gets replenished constantly. We image the entry of individual IFT motors (kinesin-II, OSM-3 and IFT-dynein) and tubulin into the cilia of the PHA/PHB neuron pair, in wild-type worms as well as in mutants lacking kinesin-2 motor and/or TZ function. Single-molecule analysis, along with numerical simulations, allows visualization of where and at what stage kinesin-2, IFT dynein and tubulin bind to assembling and/or moving anterograde IFT trains. Single-molecule localization information allows visualization of the ultrastructure and the organization at the proximal part of the cilium, also highlighting the crucial role played by kinesin-II in maintaining ciliary structure. Furthermore, we obtain a detailed picture regarding the nature of the TZ and the roles played by the two kinesin-2 motors in navigating anterograde IFT trains across the TZ.

## Results

### Single-molecule imaging reveals that kinesin-II pauses briefly before entering the cilium

To visualize the dynamics of individual ciliary proteins entering cilia, we performed SWIM on fluorescently labeled proteins in the chemosensory neurons in *C. elegans* (Mitra et al., 2022). The central idea of SWIM is to excite (and photobleach) fluorophores within only a small region of the sample (window width ∼10-15 μm), using high laser intensity, allowing continuous entry of ‘fresh’ proteins into the excited region. Here, we placed the excitation window such that we illuminated only the cilia (∼8 μm long), PCMC and small sections of the dendrites (∼1-2 μm) of a PHA/PHB neuron pair located near the tail of *C. elegans* (Figure 1A). In contrast to standard wide-field illumination, SWIM illumination allows imaging of single-molecule events for a much longer duration of time (> 20 mins; Supplementary Movie 1 and Supplementary Figure 1). In addition, the signal-to-background ratio is substantially higher using SWIM, because of the reduced out-of-focus autofluorescence background (Mitra et al., 2022).

To study the dynamics of ciliary entry, we first visualized single kinesin-II motors (using KAP-1::eGFP, non-motor subunit of heterotrimeric kinesin-2; Supplementary Movie 2; Figure 1C). We observed that kinesin-II motors show diffusive behavior in the dendrite and the PCMC, apparently not interacting with dendritic microtubules and likely in an autoinhibited state (Brunnbauer et al., 2010). At random times, individual motors appear to switch into an immobile state, close to the ciliary base (Figure 1D). Typically, they remain immobile for a while, after which they start to move along the axoneme, into the cilium, speeding up as they move past the TZ. Since anterograde IFT trains are assembled at the ciliary base (van den Hoek et al., 2022), we hypothesized that kinesin-II motors associate with or ‘dock’ to immobile, assembling IFT trains, which releases the autoinhibition of the motors, allowing them to drive the trains into the cilium. To obtain a more quantitative picture of the dynamics we extracted single-molecule tracks from the image sequences. In order to pool information from multiple cilia in different worms, we transformed the track coordinates (*x*_*i*_, *y*_*i*_) to ciliary coordinates (*c*_∥_*i*_, *c*_⊥_*i*_), by fitting a spline along the ciliary long axis and defining a reference point at the ciliary base (Figure 1E, Supplementary Figure 2 and Methods). We obtained 311 tracks of kinesin-II entry events from 12 cilia (Figure 1F). These tracks show that kinesin-II primarily docks to IFT trains between 0-400 nm from the reference point at the ciliary base, with an average docking location of 188 ± 20 nm (Figure 1G; average value and error estimated using bootstrapping). Most of the kinesin-II motors walk less than 2000 nm into the cilia, before photobleaching or detaching from the IFT train and axoneme (Prevo et al., 2015; Zhang et al., 2021). From the single-molecule tracks, we calculated that the average velocity of kinesin-II gradually increases from ∼100 nm/s at 200 nm from the start of the cilium to ∼350 nm/s at 800 nm. The velocity is relatively constant (∼350 nm/s) between 800 to 1000 nm, before increasing substantially after ∼1 μm, where the TZ ends (Figure 1H). For each track we also determined the (measured) pause time, *t*_*p*_*m*_, which we defined as the duration it takes for a kinesin-II to move 100 nm along the cilium (left panel Figure 1I). The distribution of *t*_*p*_*m*_ peaks at ∼0.75 s, with a tail extending to ∼3 s, (right panel in Figure 1I). The average pause time is 0.9 ± 0.2 s, with larger pause times (> 1.5 s) mostly observed for events docking before 400 nm from the start of the cilium (Figure 1J). We note that eGFP-labelled kinesin-II photobleaches readily and in our experiments we only sample events that have not photobleached. To better understand the effect of bleaching on our experimental observations we numerically simulated entry events, accounting for photobleaching (see Supplementary Figure 3A and Methods for details). For each simulation condition, we provided a characteristic bleach time, *t*_*bleach*_ (obtained experimentally; 1.6 s for kinesin-II; Supplementary Figure 3B), and an ‘actual’ pause time, *t*_*p*_*actual*_, and determined the measured pause time, *t*_*p*_*m*_. Parameters could be readily found (*t*_*p*_*actual*_ = 0.7 s, N=1500, N_unbleached_=320, *t*_*p*_*m*_ = 0.9 ± 0.1 s), yielding simulated distributions similar to the experimental ones (Figure 1K). The simulations reveal that the measured pause time, *t*_*p*_*m*_, increases linearly with *t*_*p*_*actual*_ before saturating at *t*_*bleach*_ (Figure 1L). Thus, our determination of pause times is affected by photobleaching when the characteristic pause time is similar to (or longer than) the characteristic bleach time of the fluorescent label. For kinesin-II, we estimate the actual pause time, *t*_*p*_*actual*_, to be 0.4-0.9 s. In summary, single-molecule analysis, complemented by numerical simulations, allows us to quantitatively describe how single kinesin-II motors associate with anterograde IFT trains, pause briefly, and enter the cilium.

### IFT dynein pauses at the ciliary base while OSM-3 and tubulin instantaneously enter the cilium

Next, we utilized this single-molecule imaging and analysis approach to understand how OSM-3 (OSM-3::mCherry), IFT dynein (XBX-1::eGFP) and tubulin dimers (TBB-4::eGFP; ciliary β-tubulin isoform (Hao et al., 2011)) enter cilia (Supplementary Movie 2). Like kinesin-II, each of these proteins appears to form a freely diffusing pool in the PCMC, with individual proteins stochastically docking to trains at the ciliary base and/or deeper inside the cilia and ultimately moving into the cilium. To our surprise, we find that the distributions of docking locations and the pause times are remarkably different for the different proteins. In comparison to kinesin-II, OSM-3 molecules attach to trains slightly deeper into the TZ (Figure 2A and 2B), with an average docking location of 334 ± 39 nm. Furthermore, as indicated by the distribution of measured pause times, *t*_*p*_*m*_ (average *t*_*p*_*m*_ 0.4 ± 0.1 s, Supplementary Figure 4A), OSM-3 molecules do not pause at all or pause much shorter than kinesin-II. Numerical simulations indicate that the actual pause time, *t*_*p*_*real*_, for OSM-3 is in the range of 0 to 0.1 s (Figure 2C; Supplementary Figure 3C). This indicates that, unlike kinesin-II, OSM-3 molecules primarily attach to already moving trains, resulting in no pause at all. In case of IFT dynein we observed that the average docking locations is 112 ± 8 nm, a very narrow region right at the ciliary base (Figure 2D & E). Furthermore, IFT dynein pauses much longer than kinesin-2 at the base, with an average *t*_*p*_*m*_ of 1.8 ± 0.2 s, and some motors pausing for more than 5 s (Supplementary Figure 4B). Numerical simulations show that the actual pause time, *t*_*p*_*actual*_, of IFT-dynein is > 9 s (Figure 2F; Supplementary Figure 3D), which is at least an order of magnitude longer than kinesin-II. For single-molecule imaging of tubulin, prolonged exposure to high intensity 491 nm laser was necessary to photobleach the bulk tubulin incorporated in the axoneme lattice. Only then ‘new’ tubulin molecules could be observed to diffuse, to move in a directed manner and/or to be incorporated into the axoneme lattice (Supplementary Movie 2). We observed that single tubulin molecules dock onto moving anterograde trains not just at the ciliary base but also throughout the TZ as well as the proximal segment of the cilium (Figure 2G and 2H; average docking location 657 ± 209 nm). Tubulin appears to dock only to moving trains, deeper inside the cilium, causing the measured (*t*_*p*_*m*_ = 0.2 ± 0 s; Supplementary Figure 4C) and the actual pause times to be negligible (*t*_*p*_*actual*_ = 0 s (Supplementary Figure 3E and Supplementary Figure 4D)). In summary, we find that tubulin, and to a lesser extent, OSM-3, dock onto already moving anterograde trains throughout the proximal part of the cilium, indicating that these proteins are also capable of diffusing through the TZ. In contrast, IFT dynein, associates with anterograde IFT trains assembling at the ciliary base, pausing for a long duration, before being carried across the TZ as passive cargo.

**Figure 2:**
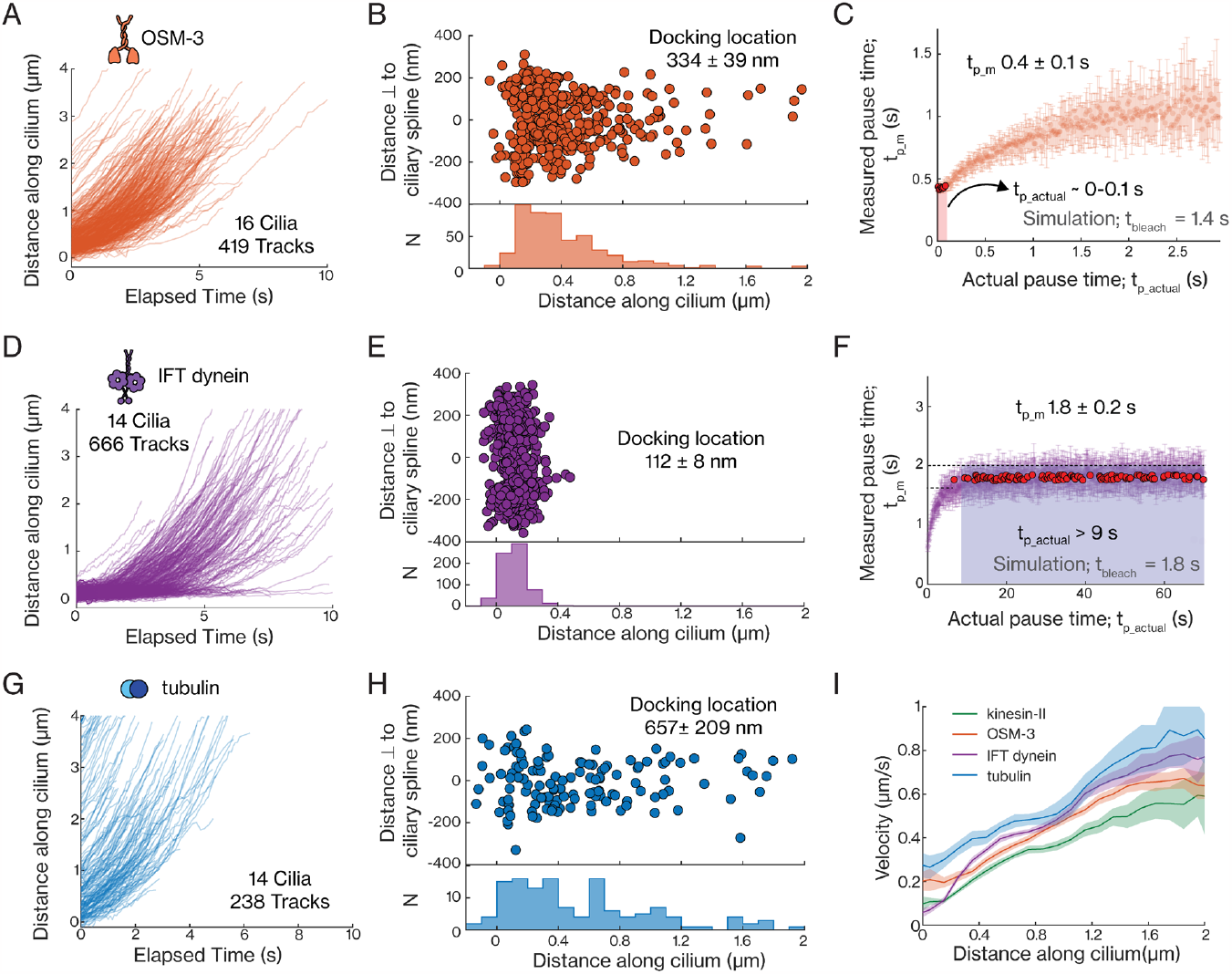
Dynamics of single OSM-3 (OSM-3::mCherry), IFT dynein (XBX-1:eGFP) and tubulin (TBB-4::eGFP) molecules entering cilia. **(A)** Distance-time plots of OSM-3 (419 tracks from 16 cilia). **(B)** Distribution (upper panel) and histogram (lower panel) of docking locations of OSM-3 (average docking location is 334 ± 39 nm). **(C)** Distribution of the measured pause time, *t*_*p*_*m*_, with respect to actual pause time, *t*_*p*_*actual*_, for OSM-3, obtained from numerical simulations (using *t*_*bleach*_ = 1.4 s). *t*_*p*_*actual*_ is in the range 0-0.1 s for *t*_*p*_*m*_ = 0.4 ± 0.1 s (experimentally obtained; Supplementary Figure 4A). **(D)** Distance-time plots of IFT dynein (666 tracks from 14 cilia). **(E)** Distribution (upper panel) and histogram (lower panel) of docking locations of IFT dynein (average docking location is 112 ± 8 nm). **(F)** Distribution of the measured pause times, *t*_*p*_*m*_, with respect to actual pause times, *t*_*p*_*actual*_, for IFT dynein, obtained from numerical simulations (using *t*_*bleach*_ = 1.8 s). *t*_*p*_*actual*_ is estimated to be > 9 s for *t*_*p*_*m*_ = 1.8 ± 0.2 s (experimentally obtained; Supplementary Figure 4B). **(G)** Distance-time plots of tubulin (238 tracks from 14 cilia). **(H)** Distribution (upper panel) and histogram (lower panel) of docking locations of tubulin (average docking location is 657 ± 209 nm). **(I)** Distribution of binned average velocities (solid line) along the length of the cilium for kinesin-II (green), OSM-3 (orange), IFT dynein (purple), tubulin (blue), with the shaded areas indicating the errors. Average values and errors are estimated using bootstrapping (see Methods).

We also looked at the average velocities of the different IFT components along the proximal part of the cilia and observed some striking differences (Figure 2I). The average velocity of OSM-3 is higher than that of kinesin-II, throughout the TZ and the proximal segment (Figure 2I). The average velocity of IFT dynein is low near the ciliary base, but increases in the TZ, becoming higher than both kinesin-II and OSM-3 (Figure 2I). Furthermore, tubulin moves significantly faster than both anterograde kinesin-2 motors all along the cilium (Figure 2I). The large variation in measured velocity of different IFT proteins can be explained in terms of differences in kinetics of docking to and unbinding from anterograde trains. It might be that kinesin-II, and to a lesser extent OSM-3, rapidly docks on and off anterograde trains in the TZ, switching into short diffusive states not detectable with the time resolution of these measurements, as reported previously (Zhang et al., 2021). This would result in a lower average velocity of single kinesin-II and OSM-3 motors. Furthermore, OSM-3 is intrinsically a substantially faster motor than kinesin-II (Pan et al., 2006). If the motor composition of IFT trains would be different for each train, trains containing more OSM-3 will be faster, resulting in a correlation between higher velocity and presence of (more) OSM-3. Finally, in contrast to both the kinesin-2 motors, it is possible that IFT dynein and tubulin, which are both cargoes of anterograde IFT trains, remain tightly coupled to the anterograde trains while they move across the TZ, resulting in the measured velocity being significantly faster.

### Single-molecule localizations allow us to map the ultrastructure of the cilium

To get insight into the molecular architecture of the ciliary base, we created super-resolution fluorescence maps of the ciliary components, making use of the single-molecule localizations in all the tracks. From all the IFT dynein tracks we obtained 30977 localizations, which we classified into ‘static’ (18561) and ‘moving’ localizations (11314) using a windowed mean squared displacement (MSD) based approach (Methods) (van Krugten et al., 2022; Zhang et al., 2021). The static localizations of IFT dynein were observed between 0 and 400 nm along the cilium, with a peak at ∼100 nm (left panel in Figure 3A) and a width of ∼500 nm (left panel in Figure 3B). This static distribution highlights the region of the ciliary base where IFT dynein docks onto assembling anterograde IFT trains. The anterograde moving IFT-dynein localizations, reveal the shape of the initial part of the cilium (right panel in Figure 3A). The distribution of localizations perpendicular to the ciliary spline is relatively broad at the ciliary base (∼500 nm wide at 0.2 μm), tapers at the TZ (∼250 nm wide between 0.4-1.2 μm) and ‘bulges’ again after the TZ, at the proximal segment of the cilia (∼400 nm wide from 1.5 μm onwards). Both for static and moving localizations, the distribution appear hollow, with fewer localizations along the ciliary spline, and more at the periphery (Figure 3B). Since the axoneme is a hollow cylinder composed of 9 microtubule doublets, we propose that the observed distribution arises from the projection of a 3D hollow cylindrical distribution undergoing a 2D projection (by our 2D imaging approach). To confirm this, we simulated localizations along the 2D cross-section of the axoneme, with input parameters, radius of the axoneme (*r*_*ax*_), radius of localizations around microtubule doublets (*r*_*db*_), and the localization error (Δ;Figure 3D). We observed that the projection of these localizations is bimodal, centred around zero (Figure 3C; Methods), similar to the experiments. We tried to estimate the underlying shape and width of the rim of the cylinder (*r*_*db*_) along which IFT components are distributed, by mapping the localization information to our simulations. We found that while we can provide meaningful information regarding the average diameter of the cylinder (Figure 3F), we cannot accurately estimate the width of the cylinder rim, due to experimental limitations, such as localization error and limited depth of field (see Supplementary Figure 5 and corresponding Supplementary text). Next, we classified the single-molecule localizations of kinesin-II and OSM-3 and obtained a negligible number of static localizations (Table 1). Super-resolution maps of moving kinesin-II (Figure 3D) and OSM-3 localizations (Figure 3E) are similar to IFT dynein (Figure 3A), suggesting that the underlying 3D distribution is the same (Figure 3G). In summary, super-resolution fluorescence maps provided by single-molecule localizations allow accurate determination of the shape of the proximal part of the cilium, showing that IFT motors move along microtubule doublets of a hollow cylindrical axoneme structure.

**Table 1:**
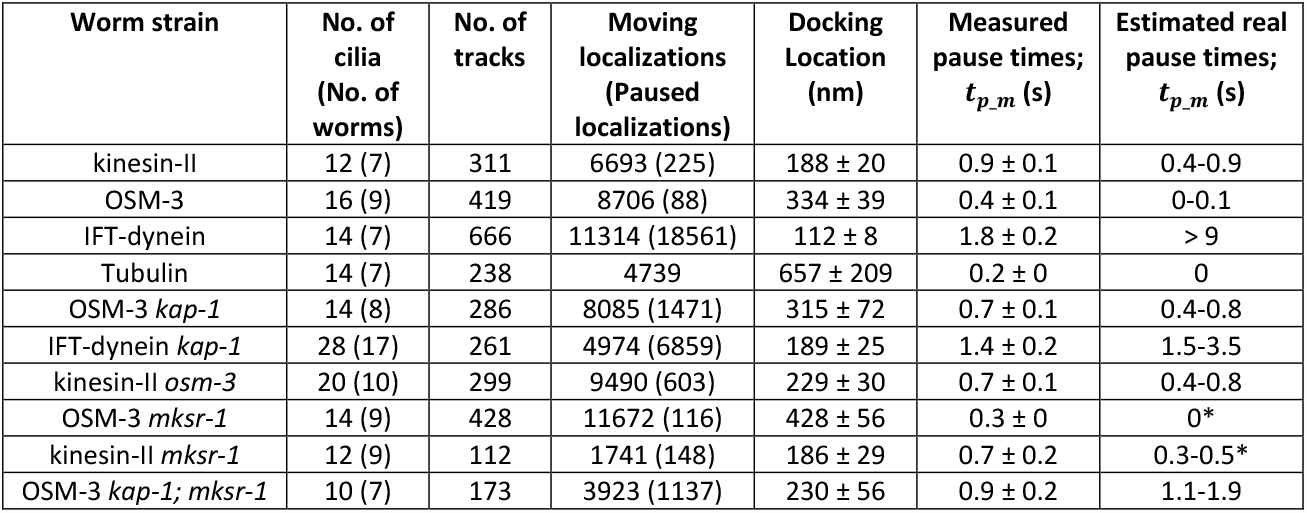
Data obtained from tracking single molecule events of different IFT components, entering the phasmid cilia. **\*** None of the simulations yield the experimentally obtained *t*_*pp*_*mm*_, hence the closest *t*_*pp*_*mm*_ is used to provide an estimate.

**Figure 3:**
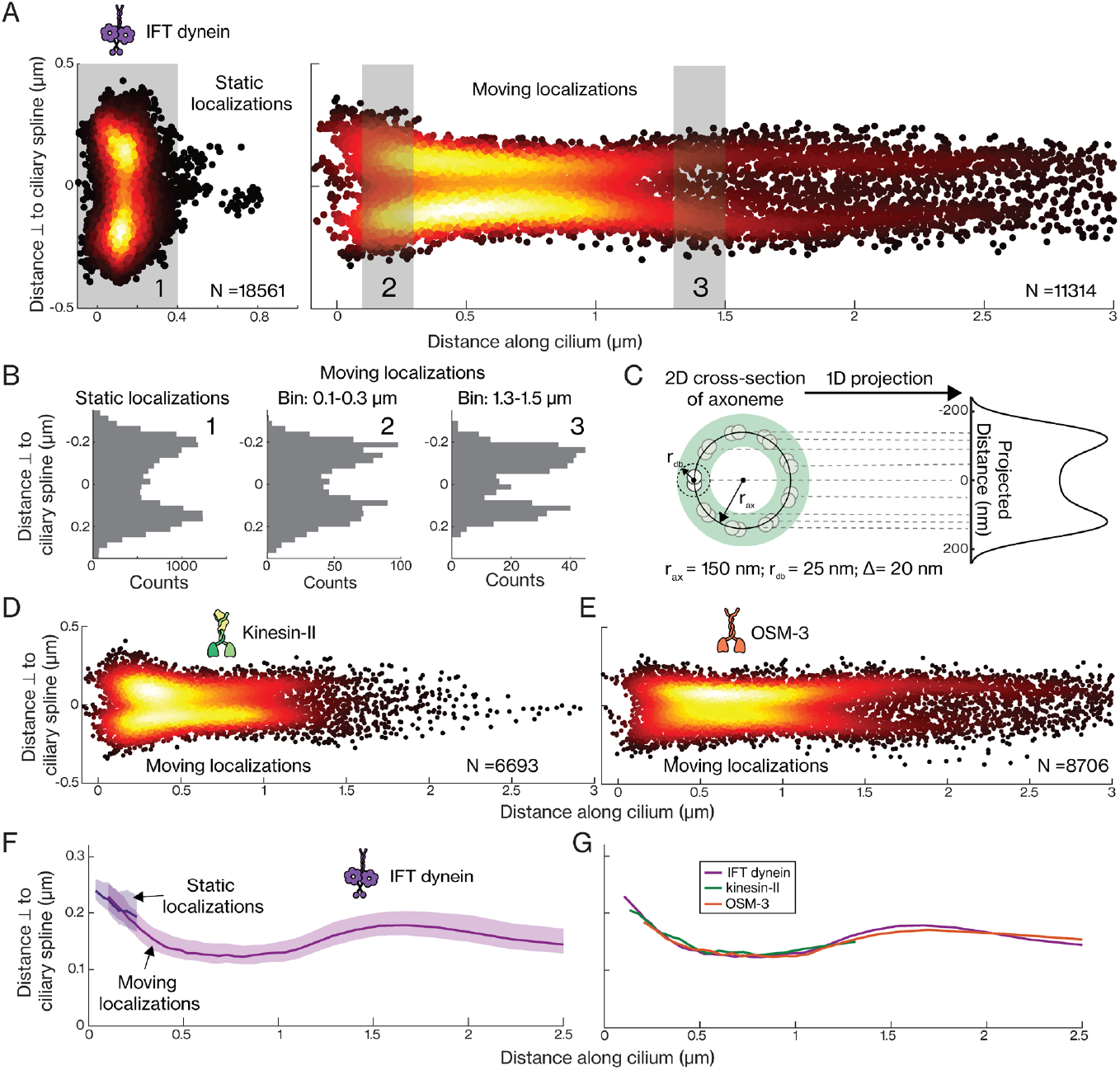
Single-molecule localizations of IFT dynein, kinesin-II and OSM-3 reveal the structure at the proximal part of cilia. **(A)** Super-resolution map of single-molecule localizations obtained from 666 IFT-dynein tracks. Left panel: N = 18561 static localizations. Right panel: N = 11314 moving localizations. **(B)** The distribution of distances perpendicular to the spline for static localizations (1), and of moving localizations between 0.1-0.3 μm (2) and between 1.3-1.5 μm (3); indicated with light grey rectangles in A. **(C)** Illustration of the transverse section of the hollow axoneme (radius *r*_*ax*_) comprising of 9 microtubule doublets, with IFT components moving along individual doublets within the radius *r*_*db*_. Right panel: 2D projection of this 3D geometry (N = 10000; *r*_*ax*_ = 175 nm, *r*_*db*_ = 20 nm and localization error Δ= 20 nm). **(D-E)** Super-resolution map of single-molecule localizations of 311 kinesin-II tracks (D; N = 6693 moving localizations) and 419 OSM-3 tracks (E; N = 8706 moving localizations). **(F)** The shape of the distribution, binned along the ciliary length, for single-molecule localizations of IFT dynein (static localizations shown in purple and moving localizations in magenta). To reflect the shape, the 80^th^ percentile value of the distribution of perpendicular distance, for every sampled bin of 1000 localizations, is plotted with the 70-90^th^ percentile range indicated by the shaded area. **(G)** The shape of the distribution, binned along the ciliary length, for single-molecule localizations of IFT dynein (magenta), kinesin-II (green), OSM-3 (orange).

### Entry dynamics of IFT components and cilia structure is altered in kinesin-II loss-of-function mutants

To obtain further insight into the roles of kinesin-II and OSM-3 at the ciliary base, we explored the entry of IFT motors in mutant worms lacking OSM-3 (*osm-3*) or kinesin-II function (*kap-1*) (Supplementary Movie 3). In the cilia of *osm-3* mutant worms, which are considerably shorter in length (Pan et al., 2006), we observed that the pausing behaviour of kinesin-II motors entering the cilium is similar to wild type (Supplementary Figure 6A and 6B), though the average velocity is slightly lower in the TZ (Supplementary Figure 6C), due to the absence of faster OSM-3 motors. In *kap-1* mutant worms, with ciliary length similar to wild type, however, the average velocity of IFT components as well as ciliary shape are very different (Oswald et al., 2018; Prevo et al., 2015). We observed that OSM-3 motors in *kap-1* worms pause significantly longer at the ciliary base (Figure 4A and Supplementary Figure 6D), with *t*_*p*_*actual*_ estimated to be in the range 0.4-0.8 s (Figure 4C). In comparison to OSM-3 in wild-type worms, the velocity of OSM-3 *kap-1* is much lower in the initial part of the TZ but increases faster, to a substantially higher velocity when the motors enter the proximal segment (Figure 4D). Docking locations show a wider distribution than in wild type, with docking occurring in the TZ as well as the ciliary base, where OSM-3 takes over the role of absent kinesin-II (Figure 4B). Remarkably, we observed that IFT-dynein motors in *kap-1* worms dock to IFT trains at the ciliary base as well as deeper inside the TZ (Figure 4E and 4F), with a velocity distribution similar to OSM-3 *kap-1* (Supplementary Figure 6E). The distribution of the measured pause time, *t*_*p*_*m*_, is also different from IFT dynein in wild-type worms, with a significantly higher fraction of IFT dynein pausing only briefly (Supplementary Figure 6F). The actual pause time, *t*_*p*_*actual*_, is estimated to be in the range 1.5-3.5 s (Supplementary Figure 6G), much shorter than in wild type. This indicates that in *kap-1* worms, either IFT-dynein motors can dock to already assembled IFT trains that are slowly moving in the TZ and/or IFT-train assembly takes less time and assembly also takes place in the initial part of the TZ.

**Figure 4:**
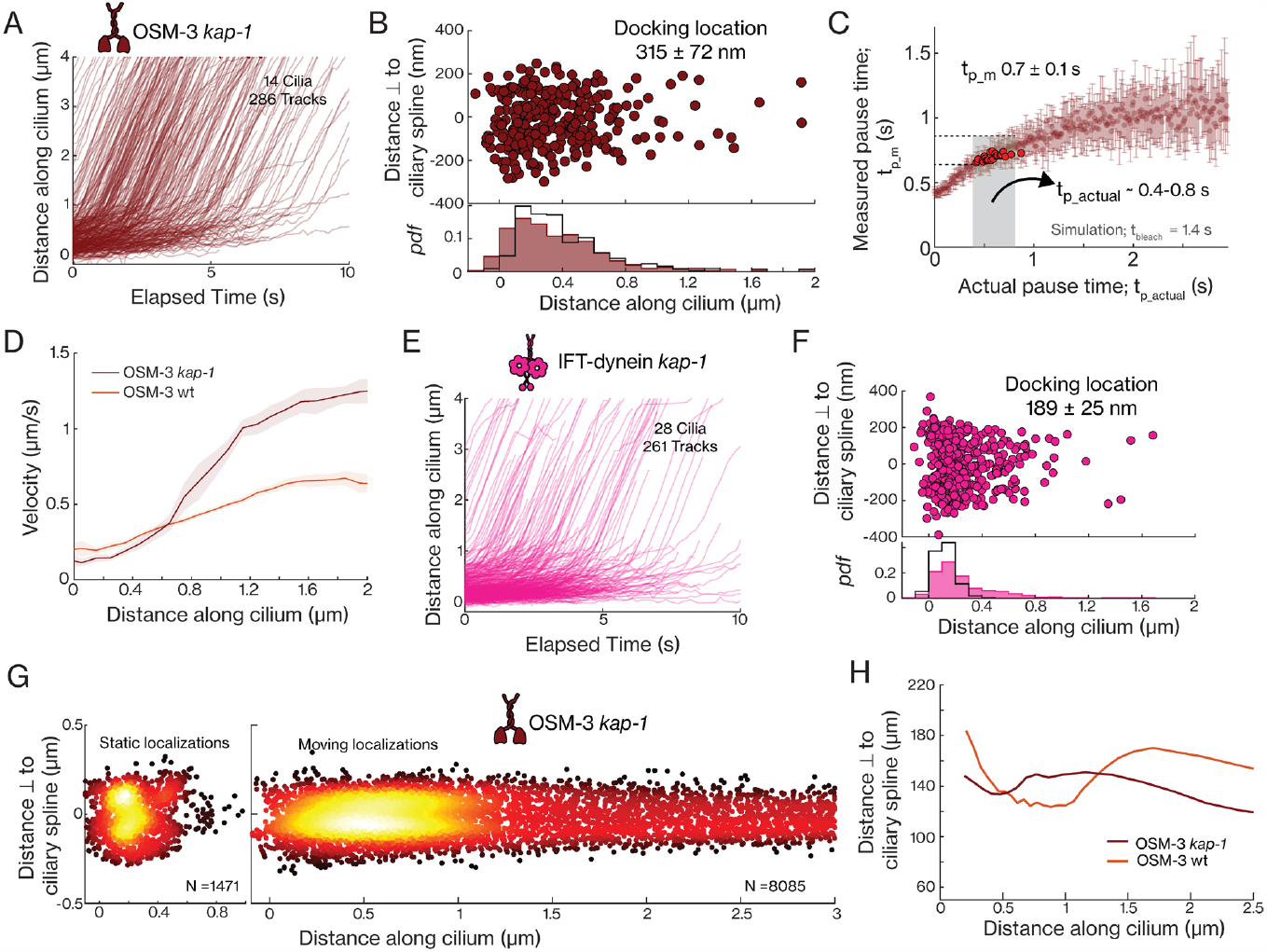
Entry dynamics of OSM-3 and IFT-dynein molecules in cilia of kinesin-II loss-of-function mutants (*kap-1*). **(A)** Distance-time plots of OSM-3 *kap-1* (286 tracks from 14 cilia). **(B)** Distribution (upper panel) and histogram (lower panel) of docking location of OSM-3 *kap-1* (brown; average docking location is 315 ± 72 nm). The distribution of docking location of wildtype OSM-3 is overlayed (black line). **(C)** Distribution of the measured pause times, *t*_*p*_*m*_, with respect to actual pause times, *t*_*p*_*actual*_, for OSM-3 *kap-1* obtained from numerical simulations (using *t*_*p*_*actual*_ = 1.4 s). *t*_*p*_*actual*_ is in the range 0.3-0.7 s for *t*_*p*_*m*_ = 0.7 ± 0.1 s (experimentally obtained; Supplementary Figure 6A) **(D)** Velocity distribution of OSM-3 along the ciliary length in wild-type (orange) and *kap-1* mutant (brown) worms. Solid line is the binned average velocity and shaded area indicates the error **(E)** Distance-time plots of IFT-dynein *kap-1* (261 tracks from 28 cilia). **(F)** Distribution (upper panel) and histogram (lower panel) of docking locations of IFT-dynein *kap-1* (pink; average docking location is 189 ± 25 nm). The distribution of docking location of wildtype IFT-dynein is overlayed (black line). **(G)** Super-resolution map of single-molecule localizations obtained from OSM-3 *kap-1* tracks. Left panel: N = 1471 paused datapoints. Right panel: N = 8085 moving datapoints. **(H)** The shape the distribution, binned along the ciliary length, for single-molecule localizations of wildtype OSM-3 (orange) and OSM-3 *kap-1* (brown). Average value and error are estimated using bootstrapping.

Super-resolution maps generated from single-molecule localizations of OSM-3 and IFT dynein in *kap-1* worms provide important insights in the ciliary structure of *kap-1* worms. First, the fraction of static localizations is significantly higher for OSM-3 in *kap-1* worms (Table 1), localizing in the initial part of the TZ, as well as at the ciliary base (left panel Figure 4G). Second, the super-resolution maps of static OSM-3 and IFT dynein show that the organization at the base is less well defined (left panel Figure 4G and left panel Supplementary Figure 6I) in comparison to wild-type cilia (left panel in Figure 3A), with static localizations distributed in more-or-less a single transversal distribution (instead of two distributions symmetrical around the centre). Third, the super-resolution maps of moving OSM-3 and IFT-dynein localizations show that the width of the cilium at the initial part of the proximal segment remains similar to the width at the TZ (Figure 4G and 4H for OSM-3 *kap-1* localizations; Supplementary Figure 6G and Supplementary Figure 6H for IFT-dynein *kap-1* localizations), while the cilium bulges at this location in wild-type worms (Figure 3G). Furthermore, the localization distributions show only one maximum in the center of the cilium (right panel Figure 4G) and not two maxima equidistant from the center, as in wild type (Figure 3A and 3E). This indicates that, in *kap-1* mutants, the axonemal structure is compromised at the proximal segment, with a shape resembling less the well-defined, hollow, 9-fold symmetric distribution of microtubule doublets and more that of randomly distributed doublets, with a less well-defined hollow space in the center. Finally, it appears that in *kap-1* mutant worms, the PHA and PHB cilia are misaligned in comparison to wild type (Supplementary Figure 6I and 6J), which might be a direct consequence of the disruption of the ciliary structure. In summary, we find that in kinesin-II loss-of-function mutants, OSM-3 can partially take over the role of kinesin-II at the ciliary base, but IFT-train assembly is affected: assembly appears to be less localized and take less time. Furthermore, the axoneme structure is disrupted, indicating that kinesin-II plays a crucial role in building and maintaining the ciliary architecture.

### IFT motors move more readily into the cilium when TZ is disrupted

In a final set of experiments, we explored the entry dynamics of kinesin-2 motors in MKS-1-related protein 1 mutant worms (*mksr-1*), in which the organization and integrity of the TZ is disrupted (Supplementary Movie 4) (Prevo et al., 2015; Williams et al., 2011). The distribution of the docking locations of OSM-3 in *mksr-1* mutant worms extends much deeper into the TZ and the proximal segment (Figure 5A and Figure 5B), suggesting that these motors can more readily enter the cilium when the TZ is compromised. The average velocities of OSM-3 (Figure 5F) and kinesin-II (Figure 5C) are higher in *mksr-1* worms, not only in the TZ, but also in the initial part of the proximal segment. In *kap-1; mksr-1* double-mutant worms, OSM-3 primarily docks at the ciliary base (Figure 5D and Figure 5E), and moves slowly through the initial part of the TZ before accelerating beyond ∼0.5 μm (Figure 5F). In comparison to the *kap-1* mutant, the velocity increase occurs earlier in the cilium and the velocity is considerably higher, even in the initial part of the proximal segment. In both *mksr-1* and *kap-1; mksr-1* worms, the higher velocity indicates that the motion of the IFT trains is less hindered and/or more kinesin-2 motors are engaged with the trains in the absence of an intact TZ, resulting in faster trains. Finally, the super-resolution map of OSM-3 in *mksr-1* worms indicates that the cilium maintains its hollow structure bulging more than in wild type (Supplementary Figure 7A and Supplementary Figure 7B). Furthermore, to our surprise, the disruption of the ciliary structure of the *kap-1* mutant (which does not show bulging of the cilium in the proximal segment) appears partially recovered in the *kap-1; mksr-1* double mutant (which shows modest bulging; Supplementary Figure 7A and Supplementary Figure 7B). Thus, we find that in TZ mutants, kinesin-2 motors enter the cilium more readily driving anterograde motion significantly faster, while the cilium maintains its structural integrity.

**Figure 5:**
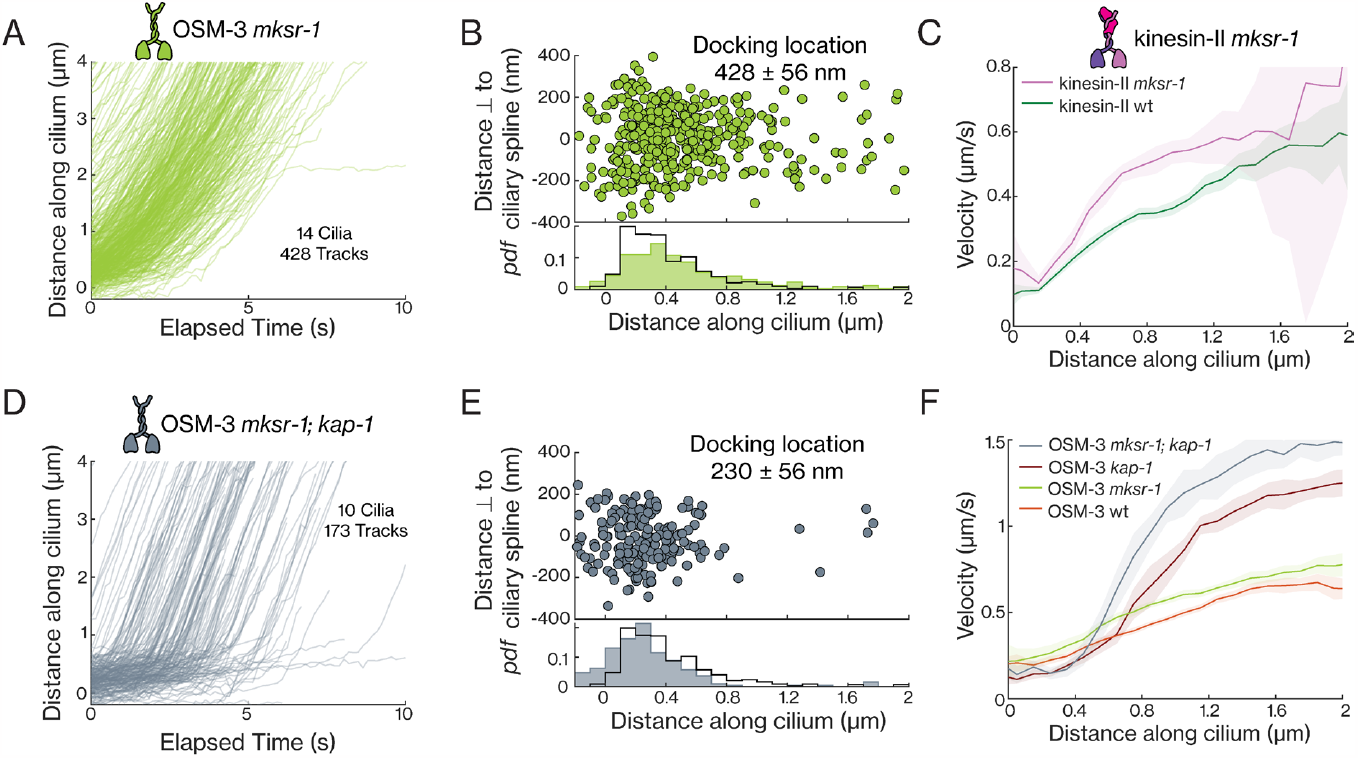
Entry dynamics of OSM-3 and kinesin-II molecules in cilia of *mksr-1* mutants and *kap-1*; *mksr-1* double mutants. **(A)** Distance-time plots of OSM-3 *mksr-1* (428 tracks from 14 cilia). **(B)** Distribution (upper panel) and histogram (lower panel) of docking location of OSM-3 *mksr-1* (average docking location is 428 ± 56 nm). The distribution of docking location of OSM-3 in wild-type worms is overlayed in black. **(C)** Velocity distribution of wild-type kinesin-II (green) and kinesin-II *mksr-1* (pink) along the ciliary length. Solid line is the binned average velocity and shaded area indicates the error. **(D)** Distance-time plots of OSM-3 *kap-1; mksr-1* (173 tracks from 10 cilia). **(E)** Histogram of docking location of OSM-3 *kap1; mksr-1* (grey; average docking location is 346 ± 44 nm). The distribution of docking locations of OSM-3 in wild-type worms is overlayed in black. **(F)** Velocity distribution of wild-type OSM-3 (orange), OSM-3 *kap-1* (brown), OSM-3 *mksr-1* (light green) and OSM-3 *kap-1; mksr-1* (grey) along the ciliary length. Solid line is the binned average velocity and shaded area indicates the error. Average value and error are estimated using bootstrapping (also in C).

## Discussion

In this study, we imaged single IFT proteins at the ciliary base of the PHA and PHB chemosensory neurons in *C. elegans* using small-window illumination microscopy (Mitra et al., 2022). We observed that the anterograde IFT motors, kinesin-II and OSM-3, as well as the cargoes of anterograde IFT trains, IFT dynein and tubulin, are present in a diffusing protein pool at the PCMC, on the dendritic side of the ciliary base. From this pool, these IFT proteins are recruited to the ciliary base by a diffusion-to-capture mechanism, as hypothesized before (Hibbard et al., 2021) and are stuck there for a while (‘dock’), before taking off on their anterograde journey through the cilium. We observed remarkable differences between the different IFT proteins. IFT dynein docks in a narrow region of space at the ciliary base and remains stuck for on average > 9 s before being transported into the cilium (Figure 2F). Kinesin-II motors remain docked for only a short while, 0.4 -0.9 s (Figure 2L), while OSM-3 and tubulin do not pause at all but start their anterograde journey immediately after docking. This points towards a sequential mechanism of anterograde IFT-train formation (Figure 6) where the core of the IFT trains is first formed by the IFT-particle complexes, to which IFT dynein binds, followed by Kinesin-II. Tubulin and OSM-3 only bind to trains that are already moving. Our findings in *C. elegans* are in line with recent studies in *C. reinhardtii*. A FRAP study showed that IFT trains take on average ∼9 s to assemble at the ciliary base (Wingfield et al., 2017) and a structural study revealed that the ciliary base is lined with anterograde IFT trains in different stages of assembly, with IFT-A and IFT dynein linearly oligomerizing from front to back on an IFT-B scaffold (van den Hoek et al., 2022). These studies and ours indicate that kinesin-2 motors can only associate with IFT trains in a late stage of their assembly. It has been shown that motor activation requires kinesins to bind to IFT trains (Brunnbauer et al., 2010; Imanishi et al., 2006), releasing autoinhibition and allowing the motors to engage with the axoneme lattice, driving the train forward. Also tubulin, which associates as cargo with IFT-B complexes in moving anterograde trains (Bhogaraju et al., 2013), does not appear to bind to the IFT trains during assembly. It is unclear what transformations an assembling train undergoes, although subtle structural differences observed between assembling and moving trains suggest that it involves a conformational change of the trains (van den Hoek et al., 2022). Further experiments will be required to unravel whether it is this conformational change that allows some IFT components to dock earlier and others later.

**Figure 6:**
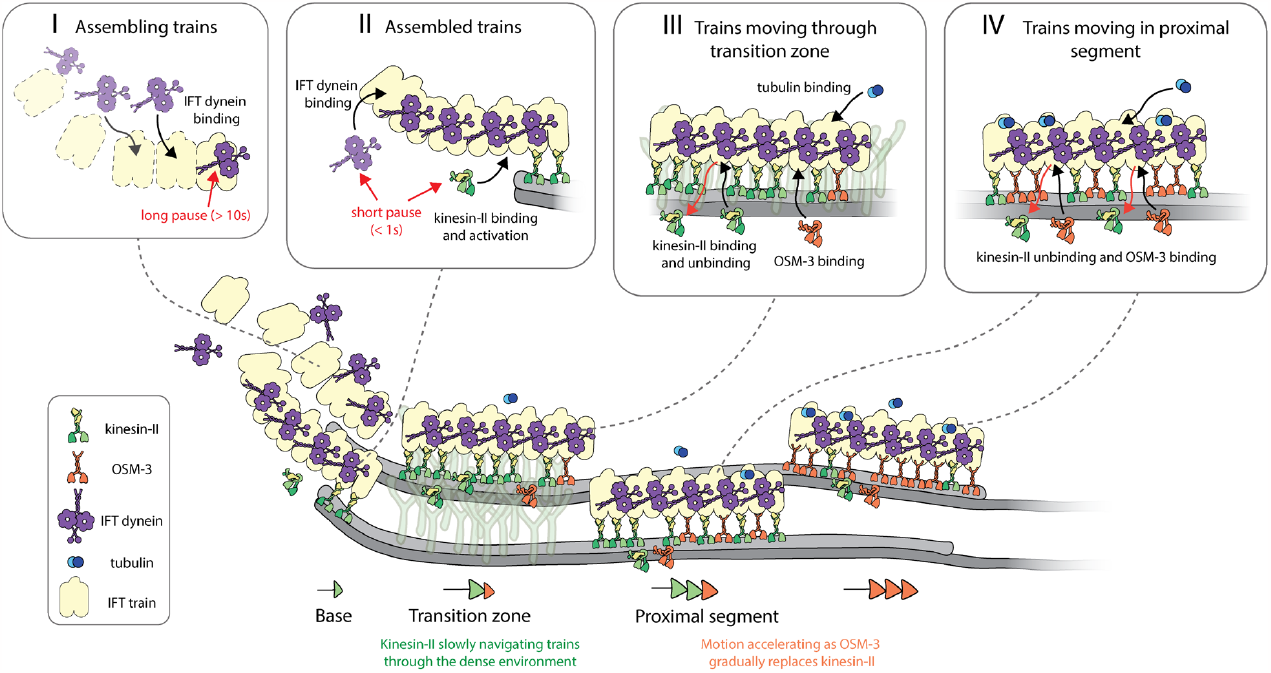
Illustration of the entry dynamics and movement of IFT motors and tubulin in cilia of *C. elegans*. Kinesin-2 motors, IFT-dynein and tubulin associate with anterograde trains at different stages. IFT-dynein molecules dock onto assembling IFT trains (see box I). When it binds to an anterograde train during the early stage of assembly, it pauses for a long duration (> 10 s). Once the anterograde IFT train is assembled, diffusing kinesin-II molecules associate with the trains, switching into an activated state (see box II). Once several kinesin-II molecules attach, the train slowly move forward. The last IFT-dynein molecules can still associate with trains in the latest stages of assembly, pausing for a short duration (< 1 s). Trains move through the TZ slowly, primarily driven by kinesin-II motors which are well suited for navigation through the dense environment (see box III). Motor (dis)association with the trains is dynamic, with kinesin-II motors detaching and both kinesin-II and OSM-3 attaching to the trains. Tubulin can also associate with the moving trains, as cargo. In the proximal segment, unhindered trains are swiftly driven by kinesin-II and OSM-3 motors (see box IV). The motion of the trains accelerates as it moves further into the cilia, since OSM-3 gradually replaces dissociating kinesin-II motors. Diffusing tubulin molecule can dock onto such fast moving trains as cargo.

In our experiments, IFT dynein only entered cilia as cargo of anterograde IFT trains (Figure 2E). We never observed IFT dynein entering a cilium by diffusing through the TZ. For tubulin, on the other hand, we observed that it associates with anterograde trains throughout the TZ as well as in the proximal segment (Figure 2G,2H), suggesting that it can freely diffuse through the TZ. This observation is consistent with studies in *C. reinhardtii*, where tubulin has been reported to enter cilia via diffusion (Craft et al., 2015; Craft Van De Weghe et al., 2020). It is well established that the TZ forms a physical boundary that prevents membrane proteins and large soluble proteins to enter cilia via diffusion (Breslow et al., 2013; Kee et al., 2012; Lin et al., 2013). In line with this, we observed that relatively small proteins like tubulin, and to a lesser extent the kinesin-2 motors, can freely diffuse over the TZ, while larger proteins like IFT-dynein and IFT trains are blocked and require IFT to cross the TZ. In *mksr-1* mutant worms, where the organization of the TZ is disrupted (Prevo et al., 2015), we observed that OSM-3 can diffuse over the TZ and associate with trains much deeper into cilia (Figure 5A, 5B), consistent with this hypothesis.

From the single-molecule trajectories we calculated the velocities of the different IFT components in the first 2-3 μm of the cilium, providing important insights into IFT in the TZ. First, we found that the velocities of all IFT components studied are significantly lower in the densely crowded TZ than further in the cilium, in the proximal segment, as has been observed before (Oswald et al., 2018; Prevo et al., 2015). Furthermore, in *mkrs-1* mutant worms with disrupted TZ, we observed that kinesin-II and OSM-3 move faster in the TZ and initial part of the proximal segment than in wild-type worms. These observations agree with previous studies showing that kinesin-driven transport slows down in a densely crowded environment (Schmidt et al., 2012; Schneider et al., 2015). Next, we found that the velocity of IFT components gradually increases in the first ∼0.6 μm of the TZ, after which the velocity remains constant. After ∼1 μm –where the TZ ends and the ciliary structure bulges out–, the velocity appears to increase again. Anterograde trains have a length of on average ∼300 nm (in *C. reinhardtii*), and consist of many IFT-B complexes, repeating every 6 nm, that provide binding sites for the kinesin-2 motors (Jordan et al., 2018; Jordan and Pigino, 2021). Most likely, in the beginning, when an IFT train takes off from the ciliary base on its anterograde journey, only a few kinesin-2 motors in the front of the train pull it forward along the axoneme, as also suggested before (van den Hoek et al., 2022). After entering the TZ, more and more motors will bind to the train and contribute to driving its anterograde transport. We hypothesize that after ∼0.6 μm into the TZ, anterograde trains have a saturating number of kinesin motors engaged with the axoneme lattice, such that trains have acquired a constant velocity in the crowded TZ. As trains move further than ∼1 μm into the cilium, they will start moving out of the TZ, resulting in less restriction by the crowded environment and further acceleration.

We also studied anterograde IFT in the absence of kinesin-II, in *kap-1* mutant worms. In these worms we observed that OSM-3-driven anterograde IFT accelerates faster beyond the TZ and that the maximum OSM-3 velocity is reached earlier in the cilium than in wild-type. This has been observed before and is due to the faster OSM-3 being the only motor responsible for anterograde transport (Mul et al., 2022; Prevo et al., 2017). In wild-type worms, the region immediately beyond the TZ coincides with the ‘handover zone’ where the slower kinesin-II starts falling off trains and more and more of the faster OSM-3 binds, resulting in a more gradual acceleration. We observed as well that the anterograde IFT velocity in the absence of kinesin-II is substantially lower in the TZ, which is in line with OSM-3 being able to take over the role of kinesin-II as importer of IFT trains and transporter in the TZ, but being less good at it. Indeed, *in vitro* studies have suggested that the slower kinesin-II is much better at navigating crowded environments than the faster OSM-3 (Brunnbauer et al., 2012; Hoeprich et al., 2014).

Another striking observation was that the velocity of kinesin-II, and to a lesser extent OSM-3, is lower than the velocities of the cargoes IFT-dynein and tubulin (∼20-30%). In a recent, higher time resolution study we have shown that kinesin-II docks on and off anterograde IFT trains frequently, switching to an autoinhibited diffusive mode after detaching from the trains (Zhang et al., 2021). The frame rate of our present experiments is likely too low to separate episodes of diffusive from processive motion. As a result, the velocity would be expected to average out to a lower value than purely processive motion. While *in vitro* studies have suggested that kinesin-II can effectively circumvent roadblocks in the crowded TZ by sidestepping on the microtubule lattice (Brunnbauer et al., 2012; Hoeprich et al., 2014), unbinding from IFT trains and rebinding after a diffusive episode could be an additional strategy to navigate in a dense environment.

The single-molecule trajectories of IFT components allowed us to generate super-resolution maps of wild-type and mutant cilia, shining further light on the functional role of kinesin-II in *C. elegans* sensory cilia. A remarkable observation was that the ciliary axoneme appears to ‘bulge’ out beyond the TZ in the proximal segment of wild-type cilia. This bulge is absent in *kap-1* mutants, as has been reported before (Oswald et al., 2018). Furthermore, in these mutants the structure of the ciliary base appeared disrupted and unorganized in comparison to wild type (Figure 4G and Supplementary Figure 6H). We also observed that IFT dynein associates with anterograde trains in a much wider region of the cilium than wild type: not only at the base, but also in the TZ and in the proximal segment. In addition, the distribution of IFT components appears less hollow in the center than in wild-type worms, which might indicate that the 9 microtubule doublets are more irregularly distributed and do not form a symmetric hollow cylinder. Taken together, these observations demonstrate that, although OSM-3 can build the cilium on its own and does not need kinesin-II, ciliary ultrastructure at the base, TZ and proximal segments appear substantially compromised in *kap-1* mutant worms. It might be that the hollow cylindrical architecture of the axoneme is necessary to properly orient trains such that they can interact with the axoneme and the ciliary membrane at the same time. In the *kap-1* strains, we also often observed that PHA and PHB cilia appear misaligned, shifted with respect to each other. It might well be that the characteristic bulge in the proximal segment of the cilia –absent in *kap*-*1*– plays a role in aligning the phasmid cilia within the channel formed by the ciliary sheath cells.

Finally, we observed that pause durations between docking of IFT-dynein to trains and departure of these trains were significantly shorter in *kap-1* mutant cilia. Before, we have measured that both frequency and size of anterograde IFT trains are lower in *kap-1* mutant cilia than in wild type (Prevo et al., 2015). Taken together, these observations indicate that, while OSM-3 is capable of taking over anterograde IFT through the TZ in *kap-1* mutants, IFT-train assembly is substantially hampered: train assembly takes shorter and trains might depart prematurely. This might indicate that kinesin-II plays a role in regulating IFT-train assembly and departure at the ciliary base. Further experiments will be needed to unravel the molecular basis of kinesin II in these processes.

In summary, we have unraveled the dynamics of individual IFT motors and cargo proteins entering the chemosensory cilia of *C. elegans*, using quantitative single-molecule imaging. Potentially, our approach can be extended to visualize the entry and exit of diverse IFT components, leading to a comprehensive understanding of the mechanisms responsible for maintaining and regulating the heterogeneous pools of proteins within the cilium.

## Methods

### *C. elegans* strains

The worm strains used in this study are listed in Table S2. The strains used have been generated before in our laboratory, using Mos-1 mediated single-copy insertion (Frokjaer-Jensen et al., 2008). Maintenance was performed using standard *C. elegans* techniques (Brenner, 1974), on NGM plates, seeded with HB101 *E. coli*.

### Fluorescence microscopy

Images were acquired using a custom-built laser-illuminated widefield fluorescence microscope, as described previously (Prevo et al., 2015; van Krugten and Peterman, 2018). Briefly, optical imaging was performed using an inverted microscope body (Nikon Ti E) with a 100x oil immersion objective (Nikon, CFI Apo TIRF 100x, N.A.: 1.49) in combination with an EMCDD camera (Andor, iXon 897) controlled using MicroManager software (v1.4). 491 nm and 561 nm DPSS lasers (Cobolt Calypso and Cobolt Jive, 50 mW) were used for laser illumination. Laser power was adjusted using an acousto-optic tuneable filter (AOTF, AA Optoelectronics). For performing small-window illumination microscopy (SWIM) (Mitra et al., 2022), the beam diameter was changed using an iris diaphragm (Thorlabs, SM1D12, ø 0.8-12 mm) mounted between the rotating diffuser and the epi lens, at a distance equal to the focal length of the latter. The full beam width in the sample was ∼30 μm (2σ of the Gaussian width). The aperture size of the diaphragm was adjusted manually to change the width of the beam, with a minimum beam width of ∼7 μm at the sample, when the diaphragm is closed to a minimum diameter of 0.8 mm. Fluorescence light was separated from the excitation light using a dichroic mirror (ZT 405/488/561; Chroma) and emission filters (525/50 and 630/92 for collecting fluorescence excited by 491 nm and 561 nm, respectively; Chroma).

For imaging live *C. elegans*, young adult hermaphrodite worms were sedated in 5mM levamisole in M9, sandwiched between an agarose pad (2% agarose in M9) and a coverslip and mounted on a microscope (van Krugten and Peterman, 2018). To perform SWIM, a small excitation window (width ranging between 10-15 μm) was used to illuminate fluorescent molecules (IFT proteins labelled with eGFP or mCherry) in a pair of PHA/PHB cilia, as well as a small section of the corresponding dendrites (see example movies in Supplementary Mov 1-4). To perform single-molecule imaging, a high intensity 491 nm or 561 nm beam (∼10 mW/mm^2^ in the centre of the beam) was used to illuminate the sample. Image acquisition was performed at the rate ranging between 6.67-20 fps, with samples typically imaged for 10-45 min. High-intensity illumination bleaches almost the whole pool of fluorescing molecules in the illuminated region of the sample, allowing visualization of ‘fresh’, not-yet bleached single molecules entering the small illuminated region.

### Image analysis

#### Tracking and spline fitting

Single-molecule events were tracked using a MATLAB-based software, FIESTA (version 1.6.0) (Ruhnow et al., 2011). Tracks, corresponding to an event, contain information regarding time (*t*_*i*_), x and y coordinates (*x*_*i*_, *y*_*i*_) and distance moved (*d*_*i*_), for every frame *i*. The connected tracks were visualized and only tracks, corresponding to single-molecule entry events starting at (or near) the ciliary base, were included for further analysis, with a minimum track length of 12 frames. All the observed IFT proteins (kinesin-II, OSM-3, IFT dynein and tubulin) diffuse in the dendrite and switch to a paused or directed state on associating with an IFT train. This switch results in a clear increase in localized intensity, marking the starting frame of the track. If diffusion was observed in a few frames before association with the IFT train, these frames were removed from the track. Erroneous tracks, primarily caused by two single-molecule events too close to discriminate, were also excluded from further analysis (though used for fitting the spline).

#### Transformation to ciliary coordinates

A ciliary coordinate system was defined by interpolating a cubic spline on a segmented line drawn along the long axis of the imaged cilium, visualized via the single-molecule localizations obtained from tracking (see Supplementary Figure 2). A reference point was picked at the base of the characteristic “bone-shaped” structure, at the ciliary base. All single-molecule localizations were transformed from x- and y-coordinates (*x*_*i*_, *y*_*i*_) to ciliary coordinates (*c*_∥_*i*_, *c*_⊥_*i*_), with *c*_∥_ the distance from the reference point along the spline and *c*_⊥_ the distance perpendicular to the spline (*c*_⊥_), as illustrated in Figure 1C.

#### Velocity measurement

Before calculating the point-to-point velocity, the tracks were smoothened by rolling frame averaging over 10 consecutive time frames, to reduce the contribution due to localization error (typically estimated to be between 10-80 nm, depending on the brightness of the tracked object). The point-to-point velocity at a given localization (*x*_*i*_, *y*_*i*_) was calculated using the following equation: *v*_*i* =_(*d*_*i*+1_ − *d*_*i*−1_)/(*t*_*i*+1_ − *t*_*i*−1_). Only moving data points (see details of classification method below) are displayed in the Figures, where we calculated the average velocity and error of the velocity data at bins of 100nm along the cilia’s long axis, using bootstrapping.

#### Docking location

Docking location of a single-molecule track was defined as the averaged coordinates (*c*_∥_, *c*_⊥_) of the first 2 frames.

#### Measured Pause time

Measured pause time, *t*_*p*_*m*_, of a single-molecule track was defined as the time taken (interpolated) to move the first 100 nm. Tracks shorter than 300 nm were discarded from the distribution.

#### Bleach time fit

The characteristic bleach time *t*_*bleach*_ for a given fluorescently-labelled protein in the cilia was obtained from the exponential fit 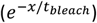 to the decay in the fluorescence intensity (normalized) over time, as a result of bleaching induced by the high intensity laser exposure (as shown for KAP-1::eGFP in Supplementary Figure 3B). The fluorescence intensity was measured on ImageJ by manually selecting a region containing fluorescent signal and a region next to it within the illuminated area, for background correction.

#### Numerical simulations of single-molecule trajectories

To determine the impact of bleaching on the measured pause time, entry events were numerically simulated (scheme illustrated in Supplementary Figure 3A). Simulated tracks were allowed to dock at a given location along a 1D cilia lattice, with the docking location randomly picked from the distribution of docking locations obtained experimentally. Each track was assigned a bleach time (*t*_*b*_) and a ‘real’ pause time (*t*_*p*_) randomly picked from exponential distributions with characteristic time *t*_*bleach*_ (obtained from experiments) and *t*_*p*_*real*_ (free parameter), respectively. The location of the simulated event was updated every Δ*t* (with Δ*t* = 150 ms, as in most experiments) and the localization precision was estimated to be 20 nm (standard deviation σ). At a frame I, while *t*_*i*_ < *t*_*b*_ or *D*_*i*_ < 6 μm (2 μm in case of kinesin-II simulation), the location of the event was updated as follows:

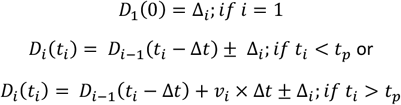

where, 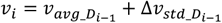 (*v*_*avg*_ is the location-dependent velocity at *D*_*i*−1_ obtained from experiments; Δ*v*_*std*_ is a randomly picked value from a normal distribution with width being the error in velocity at *D*_*i*−1_ obtained from experiments) and Δ_*i*_ is the localization precision, randomly picked from a normal distribution with width 20 nm. For each simulation condition, with number of events N, the ‘real’ pause time, *t*_*p*_*real*_, is varied and the measured pause time, *t*_*p*_*m*_, is recorded for tracks longer than 300 nm, as described for experimental data (see Figure 1H). The underlying distribution of *t*_*p*_*m*_ is estimated to be a convolution of two exponentials with rates *k*_*p*_*real*_ (1/*t*_*p*_*real*_) and *k*_*bleach*_ (1/*t*_*bleach*_) that can be defined by the following equation:

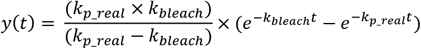

We chose to perform numerical simulations instead of fitting the data analytically since the measured pause time is different from the pause time the analytical solution provides (which cannot be measured experimentally).

#### Classification of directed transport and pausing

To obtain a quantitative measure for the directedness of the motion, we used an MSD-based approach to extract the anomalous exponent (α) from *MSD*(*τ*) = 2Γ*τ*_*α*_ (where Γ is the generalized transport coefficient and *τ* is the time lag) along the track, in the direction of motion. α is a measure of the directedness of the motion, α = 2 for purely directed motion, α = 1 for purely diffusive motion and α < 1 for sub-diffusion or pausing. For each datapoint (*c*_∥_*i*_), we calculated *α* in the direction parallel to the spline (*α*_∥_*i*_), using a windowed Mean Square Displacement classifier (wMSDc) approach, described in Danné et al. (Danné et al., 2022). α was calculated analytically, using the following equation:

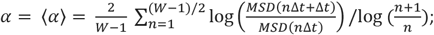

keeping a fixed window (W = 12 frames). Due to the size of the window, all tracks shorter than 12 frames were removed from the analysis. Datapoints with *α*_∥_*i*_ > 1.2 are classified as directed and *α*_∥_*i*_ < 1 are classified as static.

#### Numerical simulations to generate 1D projection library

Distribution of 2D localizations (y_*i*_, z_*i*_) along a transverse cross-section of the modelled hollow cylinder (scheme illustrated in Figure 3C), was numerically simulated (N = 10000), with *r*_*ax*_ providing the width of the cylinder, *r*_*db*_ providing the thickness of the ring and localization precision estimated to be 20 nm. Coordinates of individual localizations were obtained as follows:

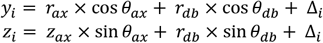

where, *θ*_*ax*_ and *θ*_*db*_was randomly assigned to the localization and Δ_*i*_ (localization precision) was randomly picked from a normal distribution with width 20 nm. The distribution of *y*_*i*_, a bimodal distribution centred at 0, was fitted with a kernel density estimation which provided the 1D projection distribution of localizations along the transverse cross-section of a hollow cylinder. A library of 1D projection distributions was generated for varying *r*_*ax*_ (ranging from 100-300 nm with steps of 5 nm) and *r*_*db*_ (ranging from 5-50 nm with steps of 1 nm). For one condition (*r*_*ax*_ = 150 nm and *r*_*db*_ = 25 nm) 1D projection distributions were obtained for varying Δ_*i*_ (ranging from 10-50 nm with steps of 10 nm; Supplementary Figure 5G). In one case (*r*_*ax*_ = 150 nm, *r*_*db*_ = 25 nm and Δ_*i*_ = 20 nm; Supplementary Figure 5H-I), a Gaussian filter of width 150 nm was applied along the z-axis, in order to under sample the localizations above and below the centre of the distribution.

#### Estimating the underlying 3D distribution from IFT-dynein single-molecule localizations

The absolute distance perpendicular to the cilia spline was sampled every 1000 data points along the long axis of the cilia (shifting every 300 data points for the next sample), from the single-molecule localization map of IFT-dynein (Supplementary Figure 5A-B). This distribution was mirrored around zero and a kernel density estimation of the distribution was obtained (Supplementary Figure 5C). By using maximum-likelihood estimation (MLE), the most accurate estimate of this distribution in the 1D projection library was determined, which allowed us to estimate the *r*_*ax*_ and *r*_*db*_.

#### Estimating the shape of the cilia from single-molecule localizations

The absolute distance perpendicular to the cilia spline was sampled every 1000 data points along the long axis of the cilia (shifting every 300 nm for the next sample) from the single-molecule localization map of a given imaged species. The 80-percentile value obtained from the cumulative distribution function of the sampled distribution provided the width of the cilia at the location along the cilia spline where the distribution was sampled. We chose the 80-percentile value of the cumulative distribution function since this is approximately where the distribution of absolute distance perpendicular to the cilia spline peaks for sampled IFT-dynein localizations. 70-90 percentile range of the cumulative distribution function was displayed to represent the error in the width.

#### Estimating average value and error for distributions

We used a bootstrapping method to calculate the parameters of a distribution (Mitra et al., 2018). We randomly selected N measurements from the distribution (with replacement) and calculated the median of the resampled group. We repeated this process 1000 times, creating a bootstrapping distribution of medians. We then calculated the mean (μ) and standard deviation (σ) of this distribution, and used these values to estimate the parameter and its error. In this paper, all values and errors are presented as μ ± 3σ.

#### Information on plots and figures

Kymographs were generated either on FIESTA or using the KymographClear (Mangeol et al., 2016) plug-in on Fiji/ImageJ. All the data was analyzed and plotted using custom written scripts on MATLAB (The Math Works, Inc., R2021a).

## Data availability

The data, and the MATLAB scripts for data analysis, visualization and numerical simulations associated with this manuscript are available on DataverseNL: https://doi.org/10.34894/EBPWXD

## Acknowledgements

We thank Dr. Misha Klein for discussions regarding the pause-time simulations and Bram Prevo for generating worm strain EJP72. We acknowledge financial support from the European Research Council under the European Union’s Horizon 2020 research and innovation programme (Grant agreement no. 788363; “HITSCIL”) and Marie Sklodowska-Curie Actions Postdoctoral Fellowship of the European Commission (Project no. 898006; ‘MingleIFT’, A.M.).

## Author contributions

Conceptualization and methodology, A.M., and E.J.G.P.; investigation, A.M. and E.L.; formal analysis, A.M.; resources, A.M.; visualization, A.M. and E.L., writing – original draft, A.M.; writing – review & editing, A.M., E.L., and E.J.G.P.; funding acquisition, A.M., E.J.G.P., supervision, E.J.G.P.

## Supplementary Information

**Supplementary Figure 1:**
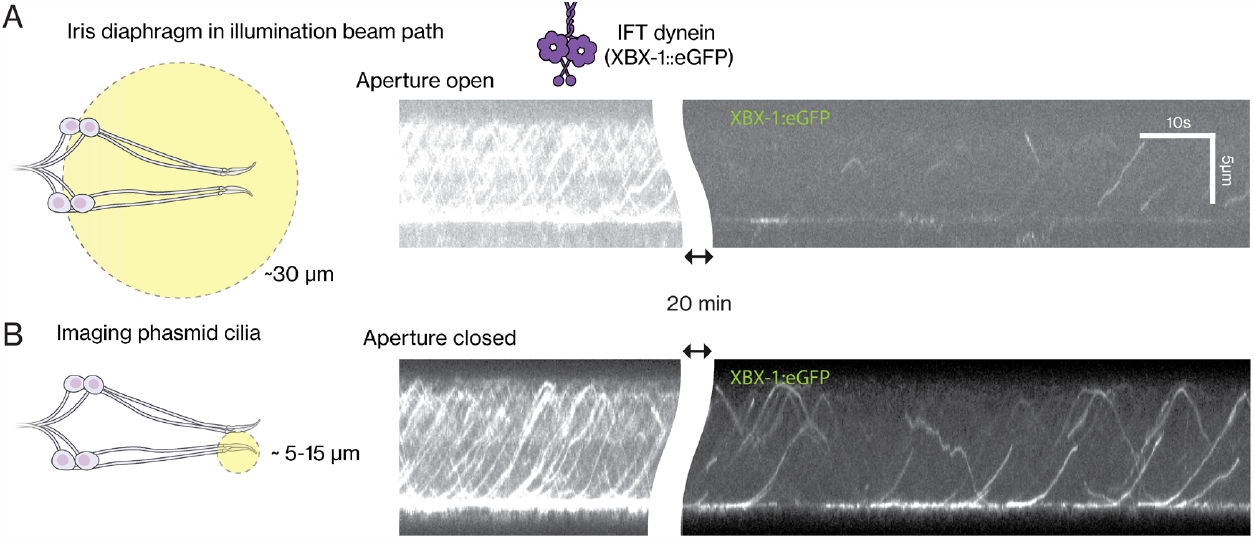
SWIM allows long duration single-molecule imaging of IFT-dynein (XBX-1::eGFP) entering the cilia. **(A)** Single-molecule imaging of IFT-dynein (XBX-1::eGFP), using the whole excitation beam (diameter ∼30 μm; aperture open) to illuminate the sample worm (illustration in left panel). Representative kymograph of IFT-dynein along an imaged cilia pair, initially shows a large pool of IFT-dynein moving back and forth in the cilia, which bleaches almost completely over time, as seen in the segment of the kymograph after 20 minutes of imaging. **(B)** Single-molecule imaging of IFT-dynein, using SWIM, exciting only a small region (diameter ∼ 5-15 μm; aperture closed; illustration in left panel). As in A, representative kymograph of IFT-dynein along an imaged cilia pair, initially shows a large pool of IFT-dynein in the cilia. After bleaching of the pool of IFT-dynein in the cilia, one can still observe unbleached IFT-dynein molecules that are continuously entering the small excited region over long duration. Since the edge of excited region is placed near the ciliary base, one can observe individual IFT-dynein molecules continuously docking at the ciliary base and entering the cilia, even after 20 minutes of imaging. Notably, the signal quality is also far superior when using SWIM. Intensity has the same scaling in the kymographs in A and B. Supplementary Movie 1 shows the movies corresponding to the kymographs.

**Supplementary Figure 2:**
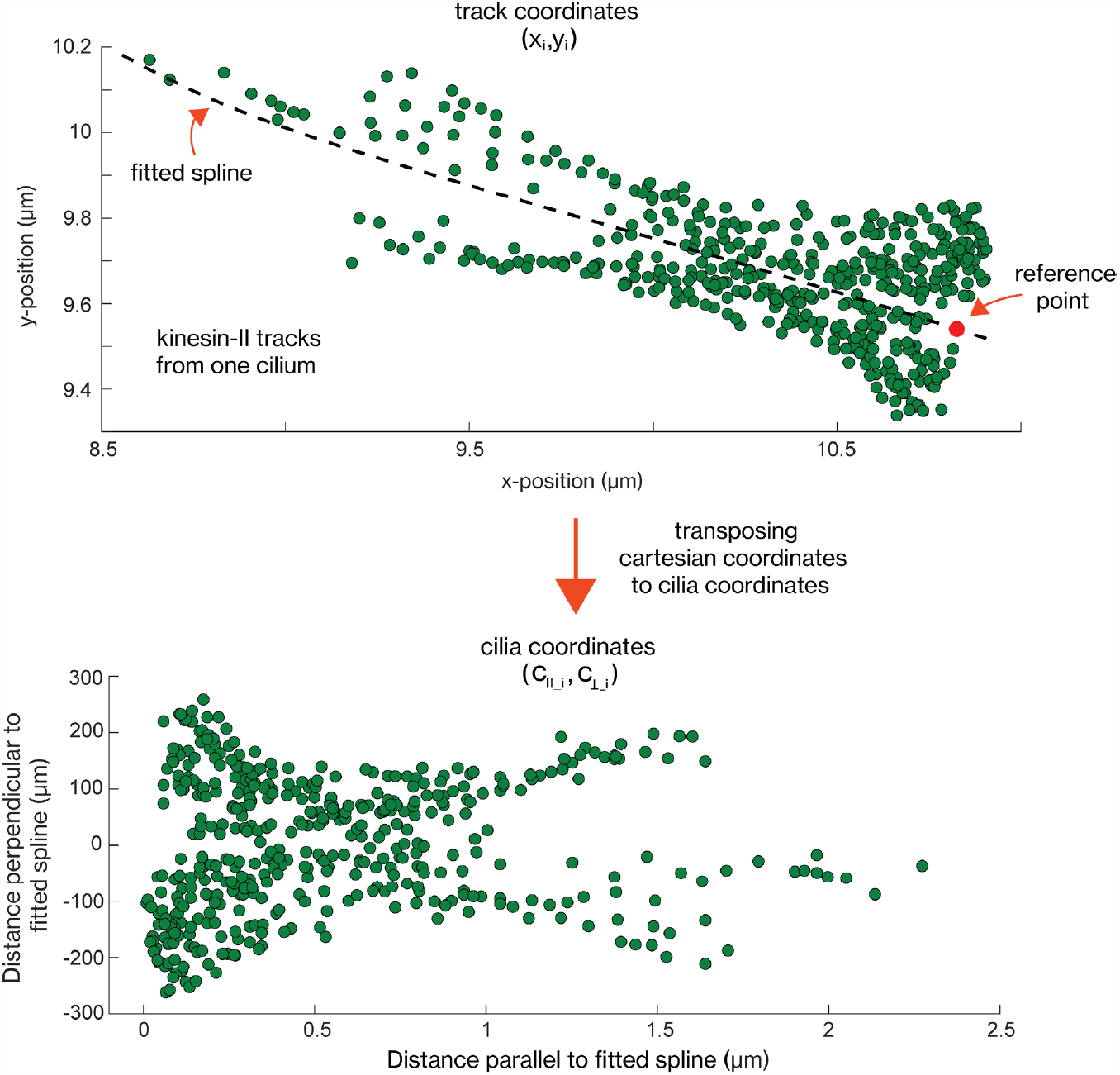
Transposing cartesian coordinates of single-molecule tracks to general cilia coordinates. Upper panel: x-y coordinates of all tracked kinesin-II entry events (476 single-molecule localizations from 20 tracks) in an imaged cilium. The single-molecule localizations provide the structure near the ciliary base, making it possible to draw a spline (dotted black line) roughly along the longitudinal axis of the cilium and select a reference point at the ciliary base. The cartesian coordinates of each single-molecule localization (*x*_*i*_, *y*_*i*_) can be transposed to distance perpendicular to the spline (*c*_⊥*i*_) and distance from the reference point (*c*_∥*i*_), referred to as cilia coordinates. Lower panel: Single-molecule localizations, plotted in cartesian coordinates in the upper panel, replotted in cilia coordinates.

**Supplementary Figure 3:**
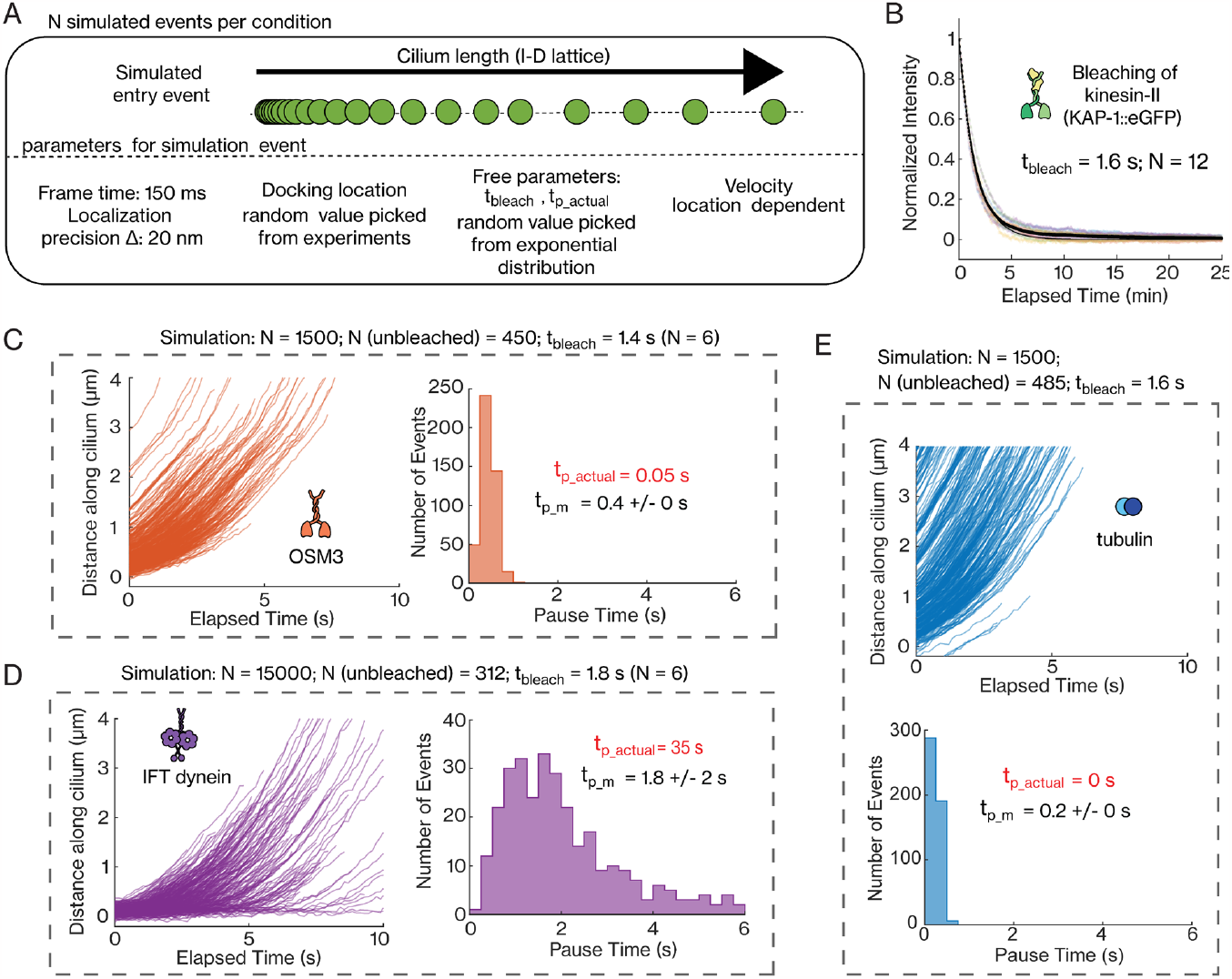
Numerical simulations to estimate actual pause time of single-molecule tracks entering cilia. **(A)** Scheme of the numerical simulation. Each simulated molecule is designated a bleach time (*t*_*b*_) and ‘actual’ pause time (*t*_*p*_), randomly picked from exponential distributions with rate parameters *t*_*bleach*_ (estimated from experiments) and *t*_*p*_*actual*_ (free parameter), respectively. The molecule docks along a 1D cilium lattice, with the docking location randomly selected from experimentally measure docking locations. After every time interval, the molecule either stays in the same location (elapsed time *t* < *t*_*b* &_ *t*_*p*_), moves forward (*t* < *t*_*b* &_ *t* > *t*_*p*_) with a location dependent velocity (obtained from experiments) or beaches (*t* ≥ *t*_*b*_; end of event). N is the number of events for a given condition, frame time is 150 ms and the localization precision is 20 nm. **(B)** Exponential decay of the kinesin-II intensity over time, upon exposure to high intensity 491 nm laser (number of cilia N = 12). The exponential fit (black line) provides a characteristic *t*_*bleach*_ = 1.6 s. **(D)** Distance-time plots of simulated OSM-3 events (N = 1500, N[unbleached] = 450) and histogram of measured pause time (average pause time *t*_*p*_*m*_ = 0.4 ± 0 s), for *t*_*p*_*actual*_ = 0.05 s. *t*_*bleach*_ is 1.4 s (N = 6). **(E)** Distance-time plots of simulated IFT-dynein events (N = 15000, N[unbleached] = 312) and histogram of measured pause time (average pause time *t*_*p*_*m*_ = 1.8 ± 0.2 s), for *t*_*p*_*actual*_ = 35 s. *t*_*bleach*_ is 1.8 s (N = 6). **(F)** Distance-time plots of simulated tubulin events (N = 1500, N[unbleached] = 485) and histogram of measured pause time (average pause time *t*_*p*_*m*_ = 0.2 ± 0 s), for *t*_*p*_*actual*_ = 0 s. *t*_*bleach*_ is 1.6 s. Average value and error are estimated using bootstrapping (see Methods).

**Supplementary Figure 4:**
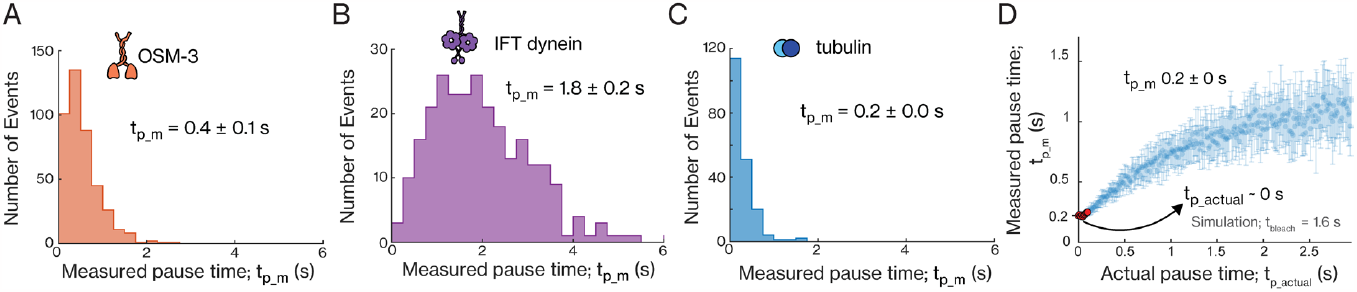
More details on dynamics of individual OSM-3, IFT-dynein and tubulin molecules entering cilia. **(A-C)** Histogram of measured pause time, *t*_*p*_*m*_, for OSM-3 (average pause time is 0.4 ± 0.1 s), IFT-dynein (average pause time is 1.8 ± 0.2 nm) and tubulin (average pause time is 0.2 ± 0 s). **(D)** Distribution of the measured pause time, *t*_*p*_*m*_, with respect to actual pause time, *t*_*p*_*actual*_, for tubulin, obtained from numerical simulations (using *t*_*bleach*_ = 1.6 s). *t*_*p*_*actual*_ is 0 s for *t*_*p*_*m*_ = 0.2 ± 0 s (experimentally obtained; 4C).

**Supplementary Figure 5:**
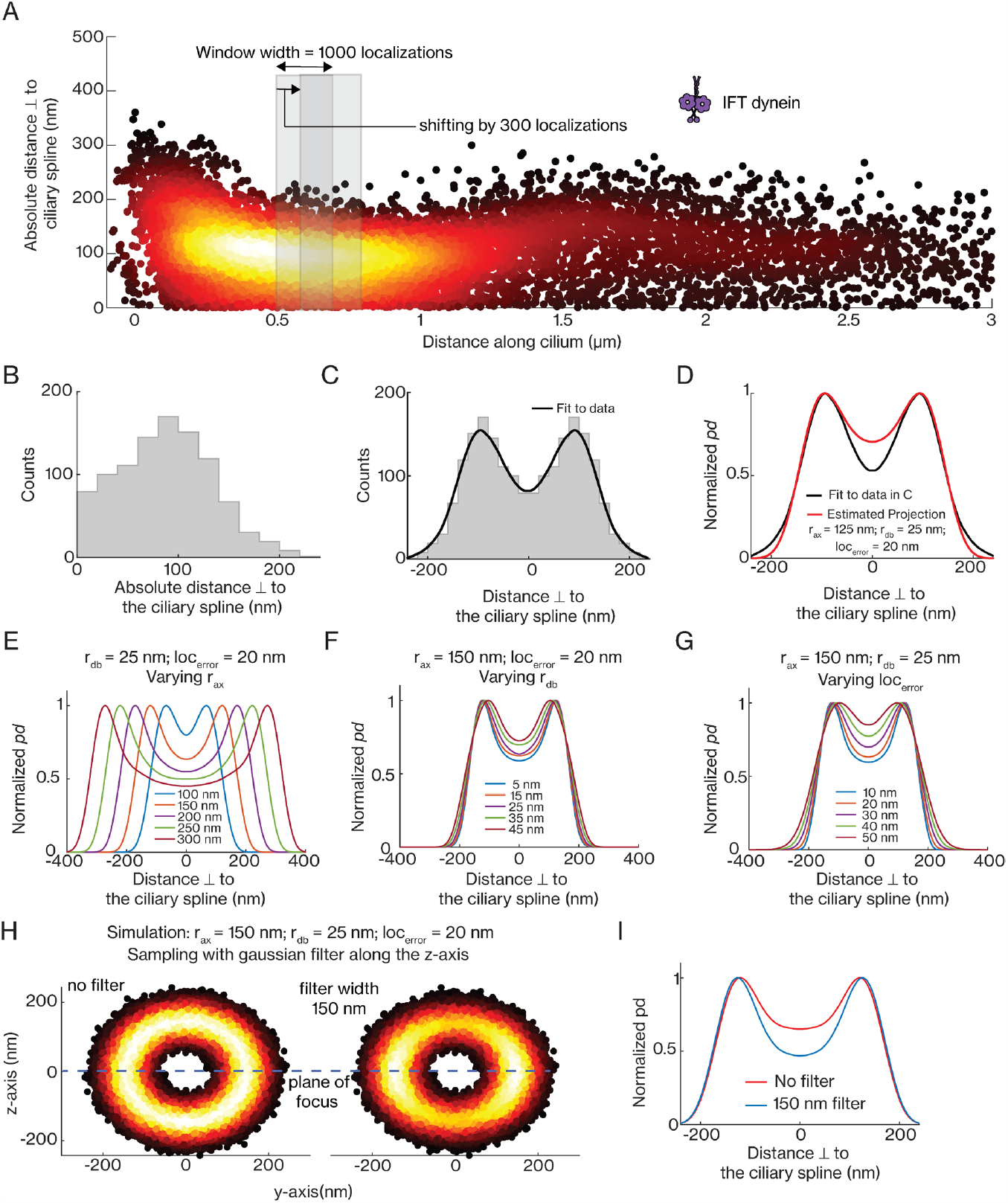
Estimating the underlying 3D distribution of the 2D projection map obtained from single-molecule localizations, using numerical simulations. **(A)** Super-resolution map of single-molecule localizations, plotting absolute distance perpendicular to the spline, obtained from 666 IFT-dynein tracks (N = 11314 moving localizations; same distribution as in left panel of Figure 3A). The distribution of distance perpendicular to the spline is sampled every 1000 data, moving every 300 data points for the next sampling. **(B)** Distribution of the absolute distance of 1000 localizations, perpendicular to the spline, located between ∼500-650 nm from the ciliary base (indicated with left shaded grey area in A). **(C)** Distribution of distance perpendicular to the spline, in B, mirrored across the spline and the kernel density fit (black line) to the distribution. **(D)** Maximum likelihood estimation of the kernel density fit in C (black), from a library of 2D projections (for a range of *r*_*ax*_ and *r*_*db*_, obtained from numerical simulations, with N = 10000 and localization error Δ = 20 nm for each condition). The best estimate for this distribution (in red) is for *r*_*ax*_ = 125 nm, *r*_*db*_ = 11 nm. **(E-G)** Distribution of 2D projections, obtained from numerical simulations (N = 10000 for each condition), varying *r*_*ax*_ (E; *r*_*db*_ = 25 nm and Δ = 20 nm; *r*_*ax*_ = 100 nm, 150 nm, 200 nm, 250 nm, 300 nm), *r*_*db*_ (F; *r*_*ax*_ = 150 nm and Δ = 20 nm; *r*_*db*_ = 5 nm, 15 nm, 25 nm, 35 nm, 45 nm) and Δ (G; *r*_*ax*_ = 150 nm and *r*_*db*_ = 25 nm; Δ = 10 nm, 20 nm, 30 nm, 40 nm, 50 nm). **(H)** Super-resolution maps of single-molecule localizations along the traverse section (yz-plane), simulated for *r*_*ax*_ = 150 nm, *r*_*db*_ = 25 nm and Δ = 20 nm (N = 10000), sampling all values (left panel; no filter) and sampling with Gaussian filter along the z-axis (right panel; σ = 150 nm for the gaussian filter). **(I)** Distribution of the 2D projections for the simulated data shown in H, sampling with (blue) and without a gaussian filter along the z-axis (red).

### Supplementary Text associated with Supplementary Figure 5

To extract the underlying structure from the experimentally obtained localizations, we generated a library of simulated 1D projections, for varying *r*_*ax*_ and *r*_*db*_ (assuming Δ to be 20 nm). We then sampled distributions of IFT-dynein localizations (distance perpendicular to cilia spline), along the cilia length (Supplementary Figure 5A; example distribution in Supplementary Figure 5B,5C) and obtained the best estimate for *r*_*ax*_ and *r*_*db*_ from the 1D projection library, using maximum likelihood estimation. We found that while the peaks of the distribution, determined by *r*_*ax*_ (Supplementary Figure 5E), can be accurately predicted, our simulations cannot fit the dip at the centre of the distribution observed in our experimental data (Supplementary Figure 5D). While we do observe that varying *r*_*db*_ (Supplementary Figure 5F) and Δ (Supplementary Figure 5G) in our simulation alters the magnitude of the dip at the centre of the distribution, the dip in the experimental distribution is much sharper. One explanation for this is that in our experiments we under-sample localizations that are above (below) the plane of focus. Indeed, when we sample simulated localizations with a Gaussian filter along the z-axis (Supplementary Figure 5H), we can replicate the distributions we observe experimentally. From this we conclude that due to experimental variables, such as localization error and under-sampling of localizations away from focus, we cannot accurately estimate the underlying 3D distributions from the 2D single-molecule localizations, in our experimental data.

**Supplementary Figure 6:**
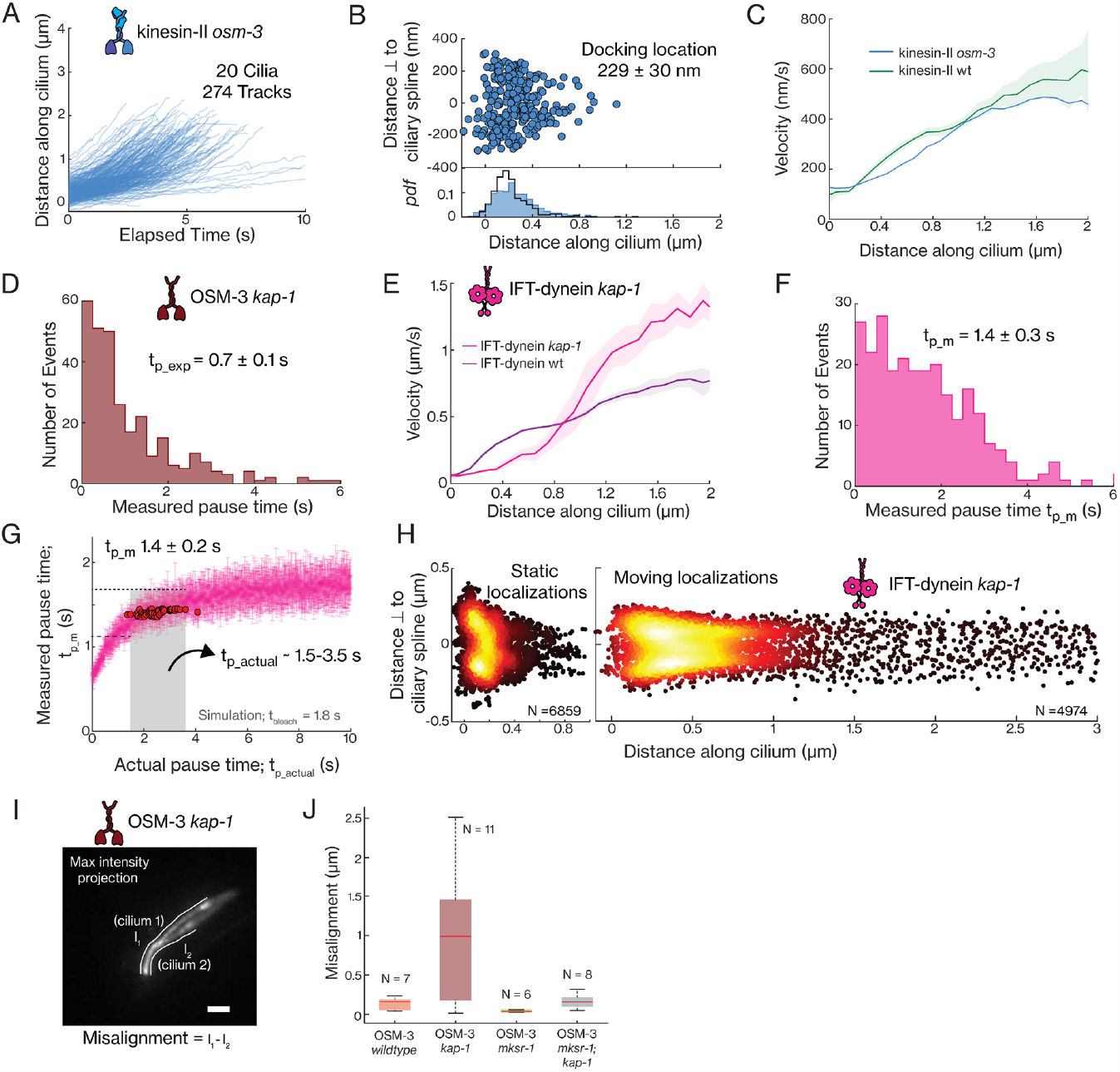
More details on entry dynamics of kinesin-II *osm-3*, OSM-3 *kap-1* and IFT-dynein *kap-1* molecules entering cilia. **(A)** Distance-time plots of kinesin-II *osm-3* (274 tracks from 20 cilia). **(B)** Distribution (upper panel) and histogram (lower panel) of docking location of kinesin-II *osm-3* (average docking location is 229 ± 30 nm). The distribution of docking location of kinesin-II in wildtype worms is overlayed in black. **(C)** Velocity distribution of kinesin-II along the ciliary length in wild type (green) and *osm-3* mutant (blue) worms. Solid line is the binned average velocity and shaded area indicates the error. **(D)** Histogram of measured pause time, *t*_*p*_*m*_, for OSM-3 *kap-1* (average pause time is 0.7 ± 0.1 s). **(E)** Velocity distribution of IFT-dynein along the ciliary length in wild type (purple) and *kap-1* mutant (pink) worms. Solid line is the binned average velocity and shaded area indicates the error. **(F)** Histogram of measured pause time, *t*_*p*_*m*_, for IFT-dynein *kap-1* (average pause time is 1.4 ± 0.3 s). **(G)** Distribution of the measured pause time, *t*_*p*_*m*_, with respect to actual pause time, *t*_*p*_*actual*_, for IFT-dynein *kap-1* obtained from numerical simulations (using *t*_*p*_*actual*_ = 1.8 s). *t*_*p*_*actual*_ is in the range 1.5-3.5 s for *t*_*p*_*m*_ = 1.4 ± 0.2 s. None of the simulations yield the experimentally obtained *t*_*p*_*m*_ (1.4 ± 0.3 s; 6C), hence the closest *t*_*p*_*m*_ is used to provide an estimate. **(H)** Super-resolution map of single-molecule localizations obtained from 261 IFT-dynein tracks. Left panel: N = 6859 static localizations. Right panel: N = 4974 moving localizations. **(I)** Time-averaged fluorescence image of OSM-3 *kap-1* shows that the cilia-pair of PHA/PHB neurons are misaligned with respect to each other. White lines of length *l*_1_ and *l*_2_ indicate the location of the cilia-pair, with the misalignment between the cilia given by *l*_1_ − *l*_2_. Scale bar is 2 μm. **(J)** Box plots of the misalignment between the two cilia in a cilia-pair, for wildtype OSM-3 (orange), OSM-3 *kap-1* (brown), OSM-3 *mksr-1* (green), OSM-3 *mksr-1; kap-1* (grey).

**Supplementary Figure 7:**
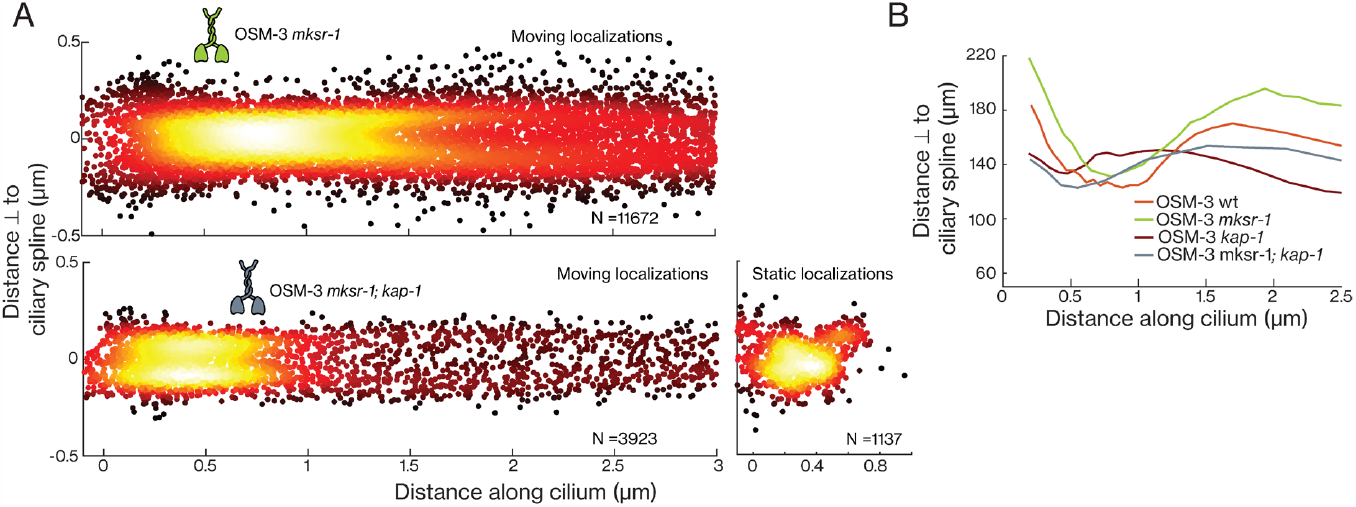
Structural details on entry dynamics of single OSM-3 *mksr-1* and OSM-3 *mksr-1; kap-1* molecules entering cilia. **(A)** Super-resolution map of single-molecule localizations obtained from 428 OSM-3 *mksr-1* and 173 OSM-3 *mksr-1; kap-1* tracks. Upper panel: N = 11672 moving localizations of OSM-3 *mskr-1*. Lower left panel: N = 3923 moving localizations of OSM-3 *mksr-1; kap-1*. Lower right panel: N = 1137 static localizations of OSM-3 *mksr-1; kap-1*. **(B)** The shape of the distribution, binned along the cilia length, for single-molecule localizations of wildtype OSM-3 (orange), OSM-3 *kap-1* (brown), OSM-3 *mksr-1* (green), OSM-3 *mksr-1; kap-1* (grey).

**Supplementary Table 1:** *C. elegans* strains used in this study. Short notation is used throughout the main text and figures to increase readability.

| Strain | Genotype | Publication | Name used |
| --- | --- | --- | --- |
| EJP13 | *Kap-1(ok676) III; vuaSi1 [pBP20; Pkap-1::kap-1::eGFP; cb-unc-119(+)] IV* | (Prevo et al., 2015) | kinesin-II |
| EJP212 | *vuaSi26 [Pxbx-1::xbx-1::EGFP; cb-unc119(+)] I; vuaSi2 [Posm-3::osm3::mCherry; cb-unc-119(+)] II; osm3(p802) IV; xbx-1(ok279) V* | (Mijalkovic et al., 2017) | IFT-dynein and OSM-3 |
| EJP401 | *vuaSi401 [pSA401; Ptbb-4::tbb-4::EGFP; cb-unc-119(+)] I; unc-119(ed3) III* | (Mijalkovic et al., 2018) | tubulin |
| EJP22 | *vuaSi2 [pBP22; Posm-3::osm-3::mCherry; cb-unc-119(+)] II; kap-1(ok676)III; osm-3(p802) IV* | (Prevo et al., 2015) | OSM-3 *kap-1* |
| EJP213 | *vuaSi26 [Pxbx-1::xbx-1::EGFP; cb-unc119(+)] I; vuaSi2 [Posm-3::osm3::mCherry; cb-unc-119(+)] II; kap1(ok676) III; osm-3(p802) IV; xbx1(ok279) V* | (Mijalkovic et al., 2017) | IFT-dynein *kap-1* |
| EJP41 | *vuaSi10 [pBP33; Pkap-1::kap-1::eGFP; cb-unc-119(+)] I; osm-3(p802) IV* | (Prevo et al., 2015) | kinesin-II *osm-3* |
| EJP72 | *vuaSi2 [pBP22; Posm-3::osm-3::mCherry; cb-unc-119(+)] II; kap-1(ok676)III; osm-3(p802) IV; mksr-1(ok2092) X* | This study | OSM-3 *mksr-1* |
| EJP64 | *Kap-1(ok676) III; vuaSi10 [pBP33; Pkap-1::kap-1::eGFP; cb-unc-119(+)] I; mksr-1(ok2092) X* | (Prevo et al., 2015) | kinesin-II *mksr-1* |
| EJP73 | *vuaSi2 [pBP22; Posm-3::osm-3::mCherry; cb-unc-119(+)] II; kap-1(ok676)III; osm-3(p802) IV; mksr-1(ok2092) X* | (Prevo et al., 2015) | OSM-3 *kap-1*; *mksr-1* |

**Supplementary Movie 1: Single-molecule imaging of IFT dynein in the cilia of *C. elegans* using the full illumination beam or using SWIM**. XBX-1::eGFP single-molecule dynamics in the cilia of PHA/PHB neurons, imaged for 25 min at the acquisition rate 6.67fps, with the aperture open (beam width ∼30 μm; left panel) and aperture closed (beam width ∼10 μm; right panel). Scale bar is 5 μm and time is indicated in min:sec.

**Supplementary Movie 2: Example single-molecule movies of IFT components in wild-type cilia**. Single-molecule dynamics of kinesin-II (KAP-1::eGFP; left panel), OSM-3 (OSM-3::mCherry; center-left panel), IFT-dynein (XBX-1::eGFP; center-right panel) and tubulin (TBB-4::eGFP; right panel) imaged using SWIM. Scale bar is 1 μm and time is indicated in min:sec.

**Supplementary Movie 3: Example single-molecule movies of IFT components in kinesin-II loss-of-function mutants**. Single-molecule dynamics of OSM-3 (OSM-3::mCherry; left panel) and IFT-dynein (XBX-1::eGFP; right panel) in *kap-1* mutant worms, imaged using SWIM. Scale bar is 5 μm and time is indicated in min:sec.

**Supplementary Movie 4: Example single-molecule movies of OSM-3 in *mksr-1* and mksr-1; kap-1 mutants**. Single-molecule dynamics of OSM-3 in mksr*-1* mutant worms (left panel) and *mksr-1; kap-1* mutant worms (right panel), imaged using SWIM. Scale bar is 5 μm and time is indicated in min:sec.

